# Decomposing neural circuit function into information processing primitives

**DOI:** 10.1101/2022.08.04.502783

**Authors:** Nicole Voges, Johannes Hausmann, Andrea Brovelli, Demian Battaglia

## Abstract

Cognitive functions arise from the coordinated activity of neural populations distributed over large-scale brain networks. However, it is challenging to understand and measure how specific aspects of neural dynamics translate into operations of information processing, and, ultimately, cognitive functions. An obstacle is that simple circuit mechanisms–such as self-sustained or propagating activity and nonlinear summation of inputs–do not directly give rise to high-level functions. Nevertheless, they already implement simple transformations of the information carried by neural activity.

Here, we propose that distinct neural circuit functions, such as stimulus representation, working memory, or selective attention stem from different combinations and types of low-level manipulations of information, or information processing primitives. To test this hypothesis, we combine approaches from information theory with computational simulations of canonical neural circuits involving one or more interacting brain regions that emulate well-defined cognitive functions. More specifically, we track the dynamics of information emergent from dynamic patterns of neural activity, using suitable quantitative metrics to detect where and when information is actively buffered (“active information storage”), transferred (“information transfer”) or non-linearly merged (“information modification”), as possible modes of low-level processing. We find that neuronal subsets maintaining representations in working memory or performing attention-related gain modulation are signaled by their boosted involvement in operations of active information storage or information modification, respectively.

Thus, information dynamics metrics, beyond detecting *which* network units participate in cognitive processing, also promise to specify *how and when* they do it, i.e., through which type of primitive computation, a capability that may be exploited for the parsing of actual experimental recordings.

## INTRODUCTION

Cognitive functions, such as working memory or selective attention, are essential for successful behavior. The associated information processing must stem from the coordinated activity of underlying neural circuits. Yet, a large gap exists between activity patterns on one side and high-level functions, which must arise through the combination of lower-level neural computations. Here, we propose a framework, capitalizing on recent advances in information theory (Lizier, 2013; Wibral et al., 2017), which aims at the data-driven detection of these simple, but yet elusive operations of raw information processing. In other words, we propose to investigate the emergence of functions arising from the dynamics of neural circuits with a specific anatomical organization by decomposing and quantifying the underlying computational building blocks.

Our proposal stems from the framework advanced by David Marr (Marr & Poggio, 1976) suggesting three possible levels of description of a neural system (cf. Figure 1A). First, “that at which the nature of a computation is expressed” (the *functional level).* Second, “that at which the algorithms that implement a computation are characterized” and “committed to particular mechanisms” (the *algorithmic level).* Third, “that at which the mechanisms are realized in hardware” (the *structural level).* Among these levels, the third, structural one is directly accessible to experimental investigation. Neural circuits across different scales have indeed a structure that can be measured (Helmstaedter et al., 2013; Binzegger et al., 2004; 2007; Angelucci et al., 2002; Markov et al., 2014) and their activity, at coarser or finer spatiotemporal resolutions, can be partially observed through a variety of recording or imaging techniques (Jercog et al., 2016; Dotson et al., 2017; Lawrence et al., 2019). Likewise, we can in many cases identify the function they ultimately give rise to, such as sensory representation, working memory, or selective attention, and measure the associated cognitive and behavioral performance, thus providing statements at the first, functional level. On the other hand, the definition and quantification of the second, algorithmic level still poses challenges. The assumption is that the algorithmic level comprises information processing operations that are intermediate steps directed toward producing meaningful functional computations but are not yet target computations. These primitive operations are pre-functional and may not have any immediately nameable goal. Yet, their alteration and disruption could lead to widespread dysfunction, since such operations would underlie disparate functional processes.

**Figure 1:**
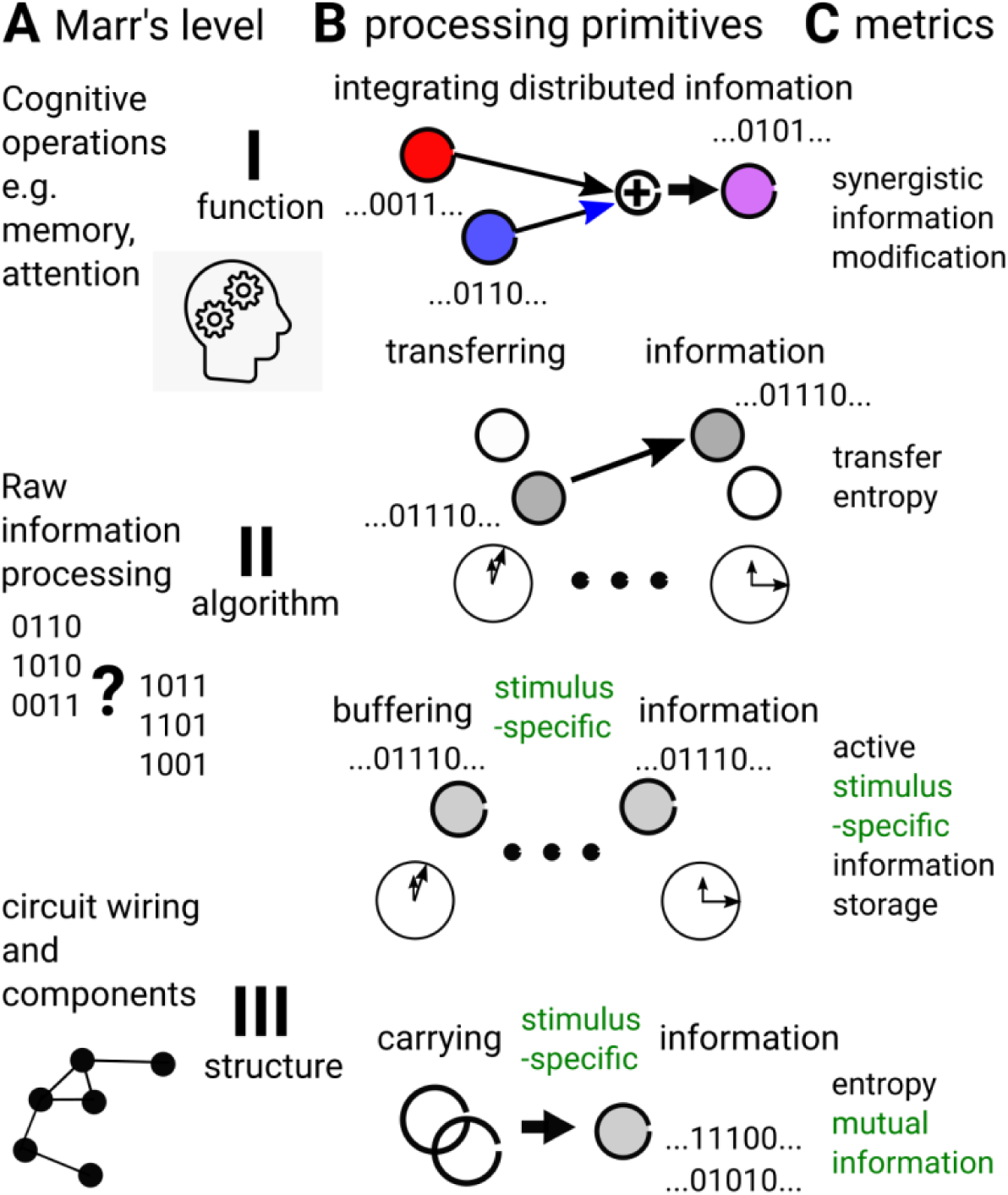
Notions of algorithmic level and information processing primitive operations. (**A**) Neural circuits can be analyzed at three different levels (Marr & Poggio, 1976): the high-level function performed by the circuit (i.e. the final cognitive operation, first *functional level*); the nature of the circuit components (neuronal types, etc.) and the anatomical wiring between them (third *structural level*); and the second *algorithmic level* of the raw information processing, bridging between circuit structure and function. (**B**) Information processing primitives (IPPs) are elementary operations performed on streams of information conveyed by neuronal activity, which are necessarily involved in different combinations in the build-up of different functions and into which more complex functions can be decomposed. (**C**) The occurrence of such IPPs can be directly tracked and quantified from neural activity data by using suitable information theoretical functionals.

Various attempts have been made to identify canonic computations that provide the building block for more complex functions. Some primarily pertained to the structural level, emphasizing the role of detailed synaptic connectivity motifs for sensory input transformations (Carandini & Heeger, 2011; Miller, 2016). Others proposed to decompose cognitive processes into simpler constituents that may be shared across multiple functions (Taatgen, 2013; Saban et al., 2021). Here, we propose a general framework that starts from the neural dynamics of neural circuits with structured and hierarchical connectivity and tries tightly linking it to emergent algorithmic operations. Indeed, neural activity conveys information and the coordination between the dynamics of interacting circuit elements eventually and unavoidably implements transformations of this information. Our goal is precisely to focus on the multitude of “informational effects” that neural dynamical patterns may induce. From the theoretical point of view, information theory precisely provides the tools to quantify the amount of information conveyed by a set of observed signals, independently from the knowledge of the meaning of this information (Shannon, 1948). Beyond assessing that certain system units convey information at a given moment in time (i.e., information is being carried “here and now”), recent developments in the Information Dynamics (Lizier, 2013) and Partial Information Decomposition (Williams & Beer, 2010; Wibral et al., 2017) frameworks also open the path to interrogate how such information is transferred and transformed, “got where it is” (i.e., to track information processing at the lowest possible algorithmic level) in an equally agnostic manner.

For instance, some fraction of the information carried by a network node’s activity may have been *transferred* to it from another location via inter-node coordination. Another fraction, may already have been present in this same node at a preceding time, thus just being actively *buffered and maintained* by the node’s activity. Another possibility is that the information carried at the current time by the node is an *integrated* assembly of pieces of information that were previously distributed in a delocalized manner across multiple nodes. Such a listing of basic operations –‘buffering’, ‘transferring’, ‘integrating’ information– does not pretend to be exhaustive. Nevertheless, it already highlights different ways to handle information and constitutes examples of possible low-level operations that we call *Information Processing Primitives* (IPPs). Classic Shannon Entropy can detect that a system’s component is ‘carrying’ information. Similarly, metrics from the Information Dynamics and Partial Information Decomposition (PID) frameworks can detect these complementary operations, with *active information storage* (Lizier et al., 2012; Wibral et al., 2014) tracking ‘buffering’, *transfer entropy* (Schreiber, 2000) tracking ‘transfer’ and propagation or *synergistic modification* (Lizier et al., 2013) tracking non-local to local ‘integration’ (cf. Figure 1B).

In the current study, we use information theoretical metrics to analyze time-series of simulated neural activity and quantify the enactment of specific IPPs. To measure which IPPs are required for the implementation of different functions–stimulus representation, working memory, selective attention, etc.–, we evaluate the proposed information-theoretical metrics on time-series produced during the execution of tasks probing the above-mentioned functions. This allows an *algorithmic decomposition* of the task, gauging the relative contribution of different IPPs without the need to reverse engineer the implementation and purpose of the performed functional computations. Such algorithmic decompositions support the intuition that different functions recruit different “flavors” of information processing, requiring varying degrees of storage, transfer or integration of information, depending on the contextual computational needs. To have a full control of every single aspect of the dynamics leading to neural function, we focus then on *computational models* of the emergence of these functions.

Simulation studies offer the advantage of arbitrarily large amounts of data in fully controlled dynamical conditions, not confounded by fluctuating brain state (Shine et al., 2016; Grossman et al., 2019), as well as a restrained noise level. Especially, the fact that such models have a known circuit architecture (third Marr’s level) and a known goal function (first Marr’s level) makes it possible to focus on the low-level algorithmic operations (second Marr’s level) bridging between the two. Our models of choice are neural circuits composed of coupled ring networks. Although stylized and analytically treatable, they retain important features of cortical connectivity, such as the spatial modulation of excitatory and inhibitory recurrent interactions. Ring networks were first introduced to study the shaping effects on feature-selective representations by collective interactions between neurons (Ben-Yishai et al., 1995) and convey an enormously rich spatiotemporal variety of dynamic patterns (Roxin et al., 2005; 2006). Multiple rings can be coupled to account for interactions between multiple cortical layers and columns (Stetter et al., 2000; Battaglia & Hansel, 2011) or even brain regions. Indeed, Ardid et al. (2007; 2010) used a network composed of two coupled rings to model the attentional modulation of responses to oriented visual stimuli. In these models, a first ring represents a cortical sensory region (e.g., region MT) producing selective responses to stimuli (e.g., oriented drifting dot patterns). These are nonlinearly gated by the interaction with a working memory copy of the same stimulus held in a second ring, representing a frontal cortical module. Here, we capitalize on these previous studies and use very similar circuit architectures, composed of one or more coupled ring networks, to perform simulated functions, such as: i) the generation of sensory responses and their maintenance in working memory; ii) the propagation of sensory responses across a cortical hierarchy of different regions; iii) the stimulus- and attention-dependent gain modulation of these responses as an effect of top-down influences. We then performed the algorithmic decomposition of these simulated functions through information-theoretical analyses of simulated tasks.

We demonstrate that data-driven information theoretical metrics are suitable to capture the different nature of Information Processing Primitives (IPPs) involved in different functions, and to identify which circuit units (neurons, populations…) are taking part in specific types of information processing in different spatial positions or moments in time. The proposed analytic framework may thus be suitable for probing the inner workings of actual cognitive processes, as they unfold during the acquisition of electrophysiological or neuroimaging recordings.

## MODEL AND METHODS

We begin with a description of the basic computational model used in this study (Figure 2) to reproduce tuned sensory responses, working memory, signal propagation through a hierarchy of areas and the modulatory effects on sensory responses of top-down selective attention. Then, we specify three setups demonstrating different types of low-level information processing underlying emulated functional computations. Finally, we detail the IPPs (i.e., the low-level information processing primitives) that characterize different types of basic information transformations. All abbreviations are summarized in Table 1 and all model parameters for the different numerical experiments performed in Figures 3–5 are specified in Table 2.

**Figure 2:**
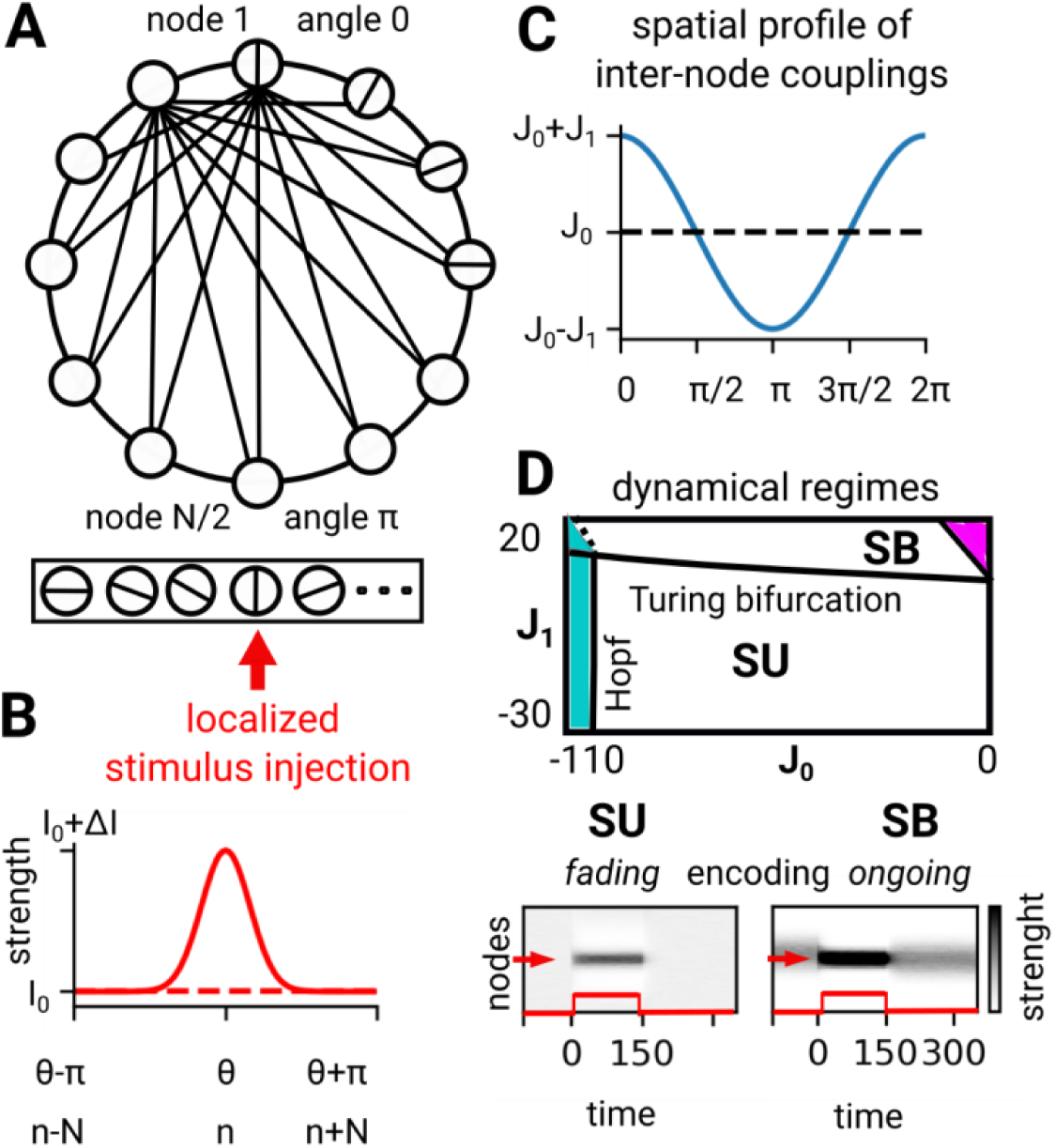
Dynamical states of the ring model of one cortical region. **(A)** Ring model to emulate information encoding and storage: circles represent feature-specific neuronal nodes, parametrized by an angle coordinate θ along the ring, indicating the preferred stimulus direction (denoted by differently oriented lines within the circles); connecting edges indicate internal all-to-all structural couplings, whose weights depend on the distance between the coupled network nodes (cf. panel **C**). The rectangle below the ring reports that stimulus-related inputs are injected in a localized fashion to network nodes with a specific stimulus-direction preference (indicated by a red arrow), following (**B**) a narrowly tuned Gaussian spatial profile. **(C)** Example profile of spatial modulation for internal ring couplings, here with a “Mexican hat” shape for parameters *J_0_*=0 and *J_1_*=1 (see *Methods*). **(D)** Top: phase diagram reporting different dynamical regimes obtained for different coupling parameter values. Bottom: Spatial maps (nodes versus time) for the two dynamical regimes explored in this study: stationary uniform (SU) activity with transient, stimulus-induced bumps of activity; and stationary bump (SB) activity with an ongoing, self-sustained bump, persisting even after stimulus offset.

**Figure 3:**
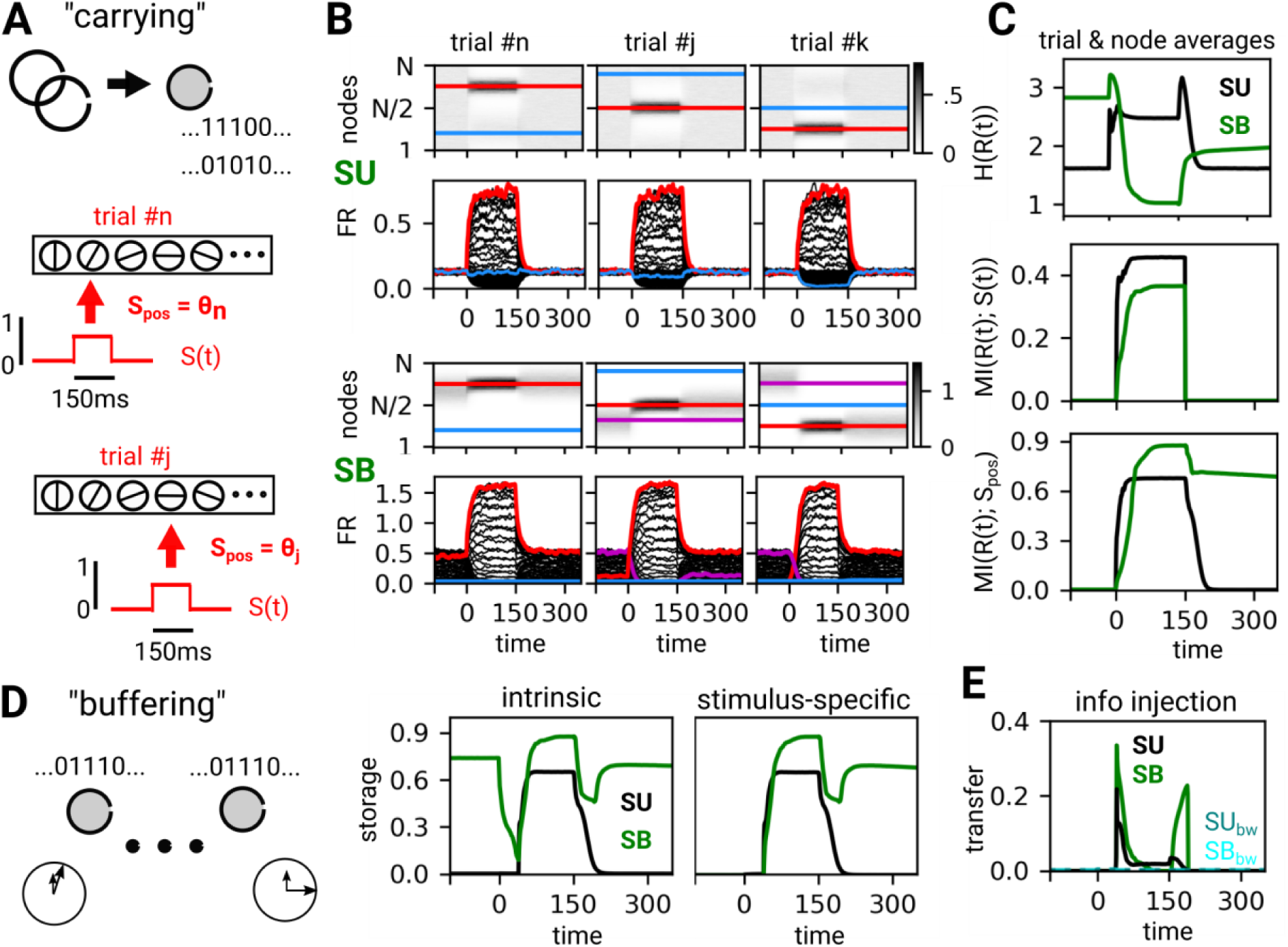
Information encoding and storage in a single-region circuit. **(A)** To track the simplest possible IPP of information “Carrying”, we simulate different trials in which stimuli with different directions θ are presented for a short, fixed time of 150 s (as indicated by the stimulus-related input time course *S(t)*). The direction *S_pos_* of the presented stimulus is denoted by a red arrow as in Fig. 1A. **(B)** Spatial maps (top row, units vs time) and single trial firing rate traces (bottom row) of neuronal activity in a one-ring network, in the SU or the SB dynamical regimes. Red lines indicate traces for nodes located at the stimulus center, blue lines nodes far from it, and magenta lines indicate the initial bump position for the SB regime. **(C)** Time-courses of entropy (top row) and stimulus-related mutual information averaged over nodes & trials (and normalized by entropy), for the SU (black) and SB (green lines) regimes. Middle and bottom row: stimulus presence and stimulus orientation are transiently encoded by activity in both the SU and SB regimes, as revealed by the mutual information between rates and, respectively *S(t)* and *S_pos_*. **(D)** Moving to the IPP of “Buffering”, we quantify and show time-courses of active information storage (intrinsic and stimulus-specific) in both the SU and SB regimes averaged over nodes & trials. Stimulus-specific storage persists after stimulus offset in SB, denoting working memory implementation. **(E)** Time-courses of information transfer from injected stimulus to rates, quantified by Transfer Entropy. Light and dark cyan lines indicate (negligible) backward transfer from rate to stimulus.

**Figure 4:**
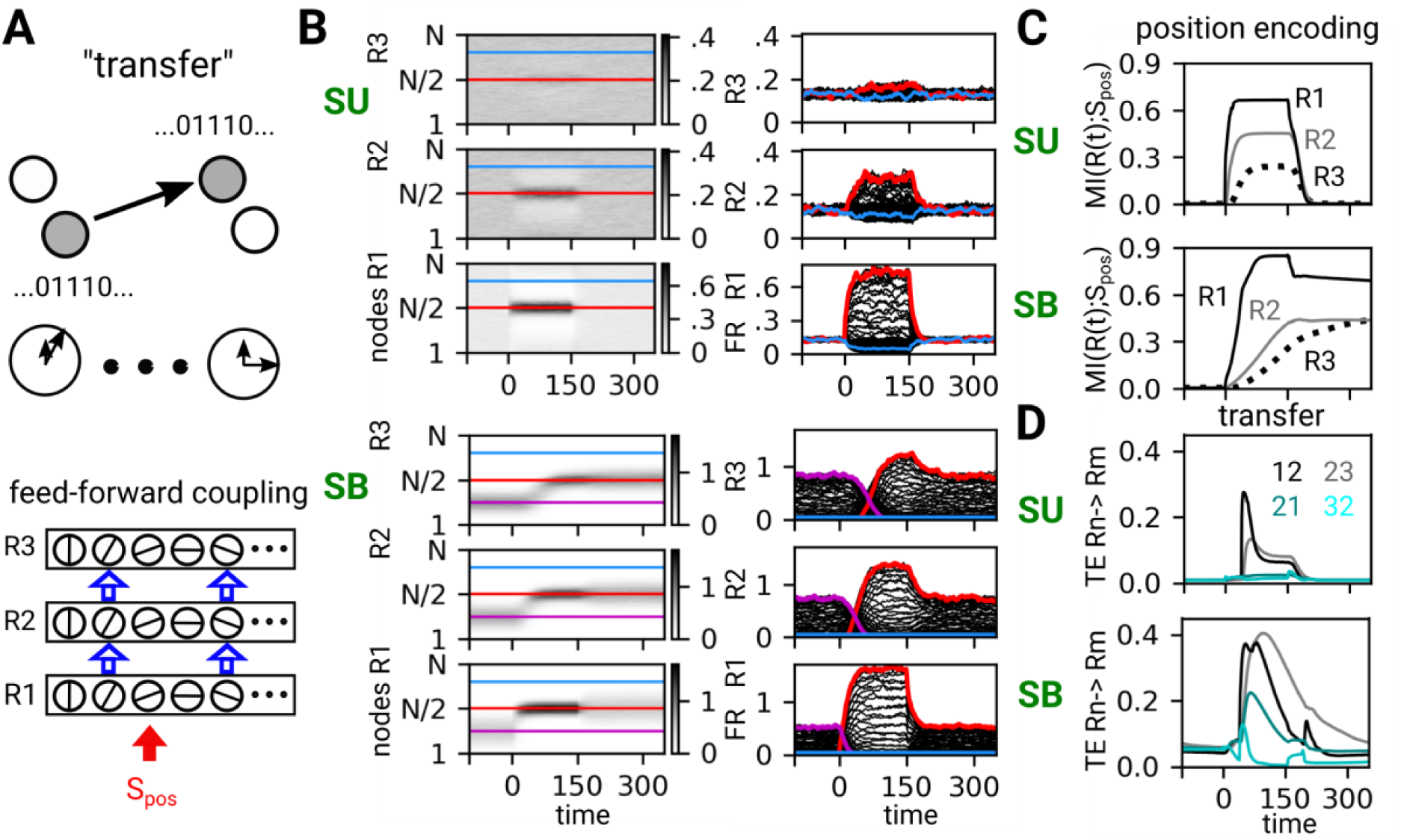
Information transfer in multi-regional feed-forward circuits. **(A)** We study the IPP of “Transferring” as it gives rise to stimulus propagation across a chain of three feed-forwarded connected regions, each modeled by a different ring network. Only the bottom ring (R1) directly receives stimulus-related inputs (red arrow). **(B)** Spatial maps (left) of single trial firing rates and corresponding rate time-series (right) in R1, R2, and R3 (top, SU; bottom, SB regimes). Red lines indicate nodes located at the stimulus center, blue lines nodes far from it, and magenta lines indicate the initial bump position for SB, as in Figure 3B. **(C)** Time-courses of relative mutual information between rates and stimulus feature (trial & node averages, entropy normalized) reveal stimulus position encoding, transient in SU (top) and persistent in SB (bottom), progressively weaker and more delayed ascending from R1 (black line) to R2 (gray line) and R3 (dotted line). **(D)** Time courses of information transfer, quantified by Transfer Entropy (trial & node averages, entropy normalized) from R1 to R2 (black), R2 to R3 (gray), R2 to R1 (dark cyan) and R3 to R2 (cyan; SU on the top, SB on the bottom). See Figure S2 for improved estimators reducing the spurious detection of backward transfer.

**Figure 5:**
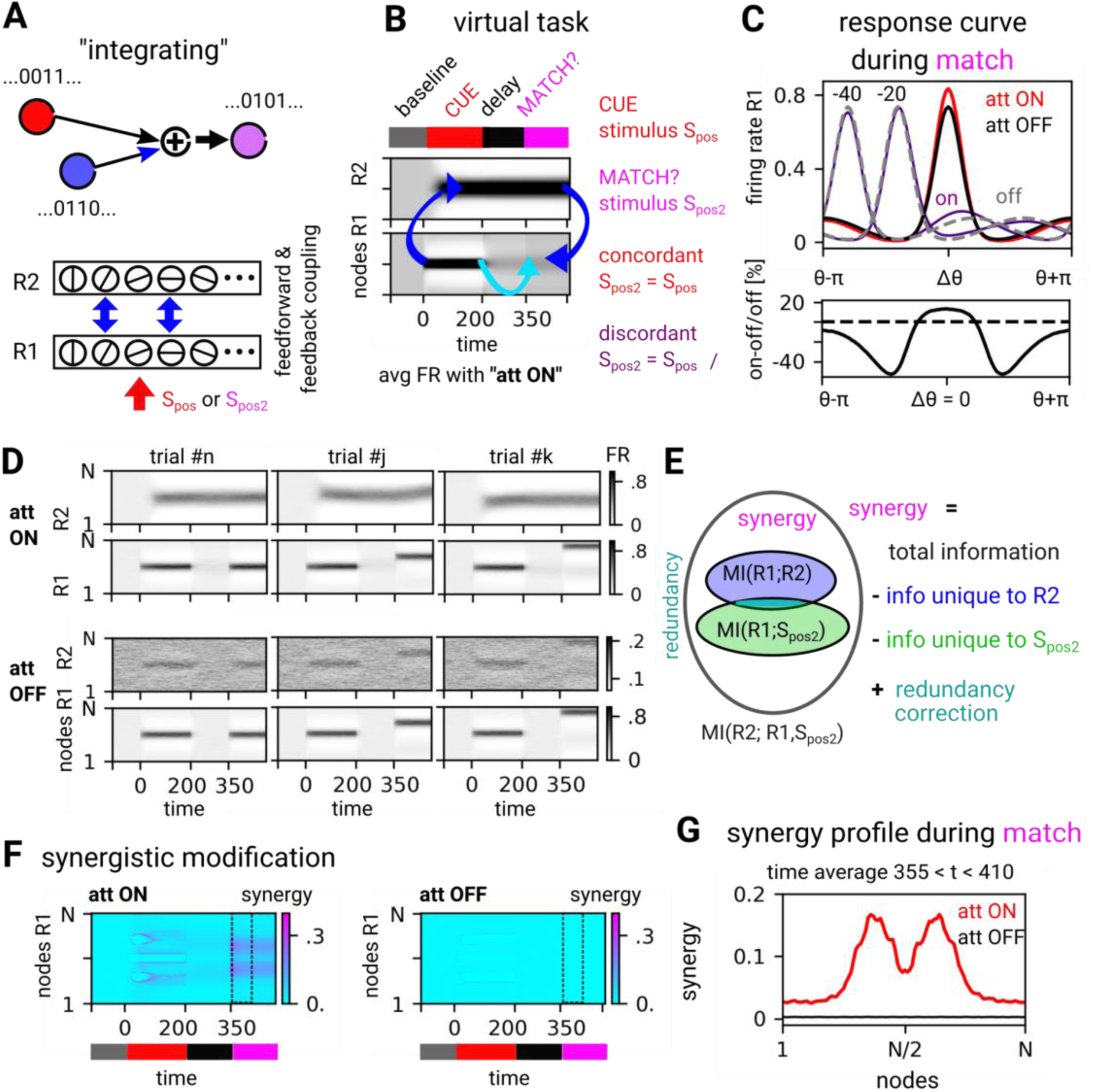
Information integration and synergy in presence of top-down attentional modulation. **(A)** We study the IPP of “Integrating” as it mediates the emergence of top-down attention-like modulation of stimulus response in a bi-regional circuit, composed of two reciprocally coupled ring networks, representing respectively a low hierarchical order sensory region and a higher-order frontal region. (**B**) In the virtual task we simulate, after a baseline period, stimuli are presented twice, during a Cue and Match stage, respectively with positions S_pos_ and S_pos2_ (red arrow), separated by a delay period without stimulation. Such a configuration mimics a selective attention experiment in which a copy of the presented stimulus is uploaded to a frontal working memory module (upward blue arrow), which stores it actively through the extent of the delay period (light blue arrow). At the moment of match, this working memory copy interacts (downward blue arrow) with the sensory representation evoked by the newly presented stimulus, matching or not the previously cued direction. During delay, the circuit can be set into an “attention ON” (upper ring in SB regime) or an “attention OFF” (upper ring in SU mode) conditions. We show here trial averaged spatial maps in the Att ON condition, in the case of matching stimuli directions (*S_pos_* = S*_pos2_*). **(C)** Response curves of firing rates averaged over trials and time during S*_pos2_* presentation (match stage). Red curve for match trials (*S_pos_* = S*_pos2_*), dotted violet curves for no-match, both for att ON. In att OFF, the black curve corresponds to a match, otherwise dotted gray curves. Bottom: attentional modulation index showing the percent enhancement (or depression) of firing rate during match stage in att ON vs att OFF conditions. **(D)** Firing rate spatial maps for three single trials with different configurations of *S_pos_*and *S_pos2_*), in bottom (R1) and top ring (R2; Top, att ON; bottom, att OFF conditions). **(E)** Venn diagram indicating the Partial Information Decomposition (PID) of the total mutual information between the sensory response in R1 and the pair of bottom-up sensory and top-down frontal inputs: Synergy equates the fraction of this total which is neither uniquely carried by R2 and S_pos2_, nor redundant between them. For other individual terms of the PID, see Figure S3. This synergistic information is extracted by nodes in R1 through the process of Information Modification. **(F)** Spatial map of the synergistic modification (normalized) in ring R1 in attention ON (left) and OFF (right) conditions. Synergy is way stronger in att ON condition, particularly in the match stage. **(G)** Section of the synergy surface during early match stage (section averaged over the time window denoted by a dotted black rectangle)

**Table 1:** List of abbreviations.

| abbreviation | full name | explanation |
| --- | --- | --- |
| SU | stationary uniform | dynamical state with spatially homogeneous activity |
| SB | stationary bump | dynamical state with spatially inhomogeneous activity |
| 3FF | three feed-forward coupled rings | setup to demonstrate information transfer |
| 2RC | two reciprocally coupled rings | setup to demonstrate information integration |
| FR | firing rates | ring model output |
| MI | mutual information | information-theoretic measure |
| TE | transfer entropy | information-theoretic measure |
| H | entropy | information-theoretic measure |
| GCMI | Gaussian copula mutual information | alternate method to calculate MI, TE, and H |
| IN | attend-IN state | configuration of the 2RC setup |
| OUT | attend-OUT state | configuration of the 2RC setup |

**Table 2:** List of parameters.

| parameter | description | values | remark |
| --- | --- | --- | --- |
| $\delta t_{int}$ | integration time step | 0.01 | arbitrary units |
| $\delta t$ | time step for the analysis | $10 \delta t_{int}$ | |
| $D$ | delay between ring units | $1 \delta t$ | |
| $D_{lr}$ | external delay between rings | $2 \delta t$ | |
| dt | time delay for MI analysis | $40 \delta t$ | for binning |
| N | number of units | 100 | 0 to 99 |
| $\nu$ | noise | 50 % | relative to the external drive |
| $J_0$ for SU state | internal coupling | -30 | stationary uniform activity |
| $J_1$ for SU state | internal coupling | -8 | stationary uniform activity |
| $J_0$ for SB state | internal coupling | -25 | stationary bump activity |
| $J_1$ for SB state | internal coupling | 11 | stationary bump activity |
| $J_2$ | external forward coupling strength | 35 | for 3FF rings |
| $\sigma_{J2}$ | external forward coupling width | 3 nodes | for 3FF rings |
| $J_2$ | external forward coupling strength | 15 | for 2RC rings |
| $\sigma_{J2}$ | external forward coupling width | 3 nodes | for 2RC rings |
| $J_3$ | external backward coupling strength | 23 | for 3FF and 2RC rings |
| $\sigma_{J3}$ | external backward coupling width | 6 nodes | for 2RC rings |
| $A_{stim}$ | constant stimulus amplitude | 2.0 | rate S(t) units |
| $\sigma_{stim}$ | stimulus width | 8 nodes | |
| $S_{pos}$ (4 features) | stimulus injection position | 0, 25, 50, 75 | for one ring and 3FF rings |
| $S_{pos}$ (1 feature) | 1st stimulus position | 50 | for 2RC rings |
| $S_{pos2}$ (10 features) | 2nd stimulus position | 0, 10... 90 | for 2RC rings |
| $t_{ON}$ | stimulus onset | $100 \delta t$ | for one ring and 3FF rings |
| $t_{OFF}$ | stimulus offset | $250 \delta t$ | for one ring and 3FF rings |
| $t_{end}$ | end of simulation | $450 \delta t$ | for one ring and 3FF rings |
| $t_{ON}$ | 1st stimulus onset | $110 \delta t$ | for 2RC rings |
| $t_{OFF}$ | 1st stimulus offset | $310 \delta t$ | for 2RC rings |
| $t_{ON2}$ | 2nd stimulus onset | $460 \delta t$ | for 2RC rings |
| $t_{end2}$ | end of simulation | $610 \delta t$ | for 2RC rings |

### Computational model of one region: ring network model

The building block of all networks studied here is the rate-model version of the one-dimensional (1D) ring network with delayed interactions described by Roxin et al. (2005; 2006), expanding on a classic model by Ben-Yishai et al. (1995), see Fig. 2A. The one-ring network provides a canonical model for a feature-selective cortical module (e.g., a visual cortex hypercolumn). In ring rate models, the activity of *N* coupled network nodes (also called units) is characterized by their firing rates *R_k_(t)*, *k* = 0…*N*-1. The total input to each unit *k* is a linear combination of the activity of all presynaptic units, an external stimulus, and an external drive *I_ext_*, set to have a baseline stationary rate equal to *R_k_(t)* = 0.1.

Each unit may receive at certain times additional external input *I_stim_*, associated with the presentation of an external stimulus. Such stimulus current is spatially localized, to model the stimulus selectivity of different nodes. We choose the form of a Gaussian kernel (cf. Fig. 2B) of prescribed maximum amplitude *A_stim_* and width σ*_stim_* (see next section for the rationale on their selection), centered at a position 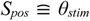, which varies across different simulated trials. The time-course of the stimulus is given by a function *S(t),* equal to one during stimulus presentation and zero otherwise.

The time evolution of the activity of each node is governed by a first-order delay differential equation with time delay *D* involving a threshold-linear input-output transfer function *Φ(x) = x* if *x* > 0 and *Φ(x)* = 0 otherwise:

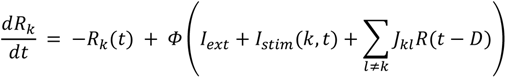

We consider ring modules (i.e., regions) of *N* = 100 nodes, where each node *k* is labeled by its angular position on the ring *θ_k_* = 2π*k*/*N*, *k* = 0, 1 . . *N*-1, coupled to all other nodes *l* ≠ *k* through a distance-dependent coupling kernel *J_kl_*, depending on the angular distance between nodes:

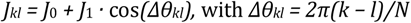

for the link between nodes *k* and *l*. The coefficients *J*_0_ and *J*_1_ control the spatial modulation and crucially, the net sign of interactions (excitatory or inhibitory) between rate units. Fig. 2C shows an example of coupling kernel for the parameter choices *J*_0_ = 0 and *J*_1_ = 1, resulting in excitatory short-range interactions with nodes within a range −π/2 ≤ Δ*θ_kl_* ≤ π/2, and in inhibitory long-range interactions with nodes farther away, i.e. |Δ*θ_kl_*| > π/2 (“Mexican hat” profile).

### Dynamical regimes of the ring model and properties of stimulus response

The ring network exhibits a rich spectrum of dynamical states, as a function of the parameters *J_0_* and *J_1_*. Fig. 2D (top) shows a schematic phase diagram (exemplary with noise strength of 50% baseline external drive). As studied in detail by Roxin et al. (2005; 2006), the model exhibits a stationary fixed-point solution for small coupling values. This corresponds to an asynchronous regime of firing in which the average firing rate is constant in time and spatially homogeneous throughout the ring in absence of external stimuli (*Stationary Uniform regime*, *SU*). When modifying the *J*_0_ and *J*_1_ parameters, the SU regime loses stability. For large *J*_0_ or *J*_1_ the firing rate of every unit explodes toward infinitely large values (“rate instability”), as the chosen threshold-linear transfer function does not saturate. However, when *J*_0_ < 0, i.e. collective interactions are negative on average, there is a finite range of positive *J*_1_ Mexican-hat modulation, for which the system’s activity does not explode, but spontaneously gives rise to localized bumps of activity. These are centered at some stochastically selected angular position (spontaneous symmetry breaking or “Turing instability”) and surrounded by silent units (*Stationary Bump regime, SB*). Finally, when the average interaction level becomes strongly inhibitory for *J_0_* << 0, the SU regime undergoes a transition (“Hopf instability”) to a regime in which the firing rate oscillates homogeneously and in-phase throughout the ring at a finite frequency. This *Oscillatory Uniform regime (OU)*, together with additional regimes (such as traveling waves, etc.) that the delayed ring-models could express in other parameter ranges (Roxin et al., 2005; 2006), is not further explored in this study.

We focus on the SU and SB regimes, notably on their responses to externally presented stimuli (Fig. 2D, bottom). In absence of a stimulus, the activation in the SU regime is uniform throughout the network, as previously described. However, when a stimulus is presented at a certain angle *θ_stim_*, then a bump of stronger activity, centered on this angle *θ_stim_*, develops due to the locally increased excitatory drive. This, in turn, also silences the surrounding nodes outside of the bump via lateral inhibition. Such a bump can be seen as a representation of the presented stimulus, as its position along the ring follows the angle of the external stimulus. In the SU regime, when the stimulus presentation ends, i.e. the additional *I_stim_* input goes back to zero, the bumps dissolve and activity relaxes back to uniform (fading encoding, Fig. 2D bottom left).

The situation is different in the SB regime where a spontaneously generated bump is already present before the stimulus occurs. In this case, the effect of presenting an external oriented stimulus is not the creation of a bump, but rather the displacement of the previously existing bump, moving it to the location corresponding to the presented stimulus angle *θ_stim_*. Once the stimulus is removed, the evoked bump continues to exist, because it is self-sustained by local recurrent excitation. It will persist for a certain time at the new position (persistent encoding, Fig. 2D bottom right), before noise causes it to drift. In the following, we perform simulations at selected working points within the SU and the SB regimes (see Table 2 for our parameters choices). Note that the parameters *A_stim_* and σ*_stim_* of the stimulus are chosen and tuned in such a way that the bump response evoked by a stimulus in said SU working point has similar width and amplitude to the bump arising in the SB working point, such that the results obtained from simulations in the two regimes are more easily comparable.

### Multi-regional architectures: coupled ring networks

To model circuits involving more cortical modules (e.g., different stimulus-representing regions), we use networks composed of multiple coupled rings. In these multi-ring architectures, each regional module (i.e., ring) is modeled as the previously described single-ring architecture (with the possibility to tune different regions to different dynamical regimes and working points). However, an additional current term *I_lr_* must be fed to the transfer function Φ of each unit *R_k_* to account for an additional drive provided by the long-range (lr) coupling to nodes in remote rings:

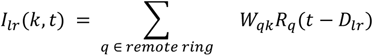

The kernel *W_rqk_* of long-range connectivity, for simplicity, has a rectangular shape. It assumes a positive amplitude *A_lr_*within the range *k*-σ*_lr_* ≤ *q ≤ k+*σ*_lr_* (thus symmetrically centered on *k*), and zero outside this range, ensuring a strict spatiotopy of inter-regional excitatory projections. Inter-ring interactions are also delayed, analogous to the recurrent interactions within rings, and can have an independently tuned, longer delay *D_lr_*. We here consider two types of multi-ring architectures.

*Three-rings network with feed-forward coupling.* After exploring representation and maintenance of a stimulus representation by a single sensory region (Figure 3), we study the propagation of such a representation through a hierarchy of different sensory areas (Figure 4). To emulate the transfer of information from one area to another, we couple three rings as a feed-forward chain (Fig. 4A). We refer to this setup with the label “3FF rings” (cf. Tables 1 and 2). The bottom ring (R1) represents a sensory cortical area which directly receives subcortical stimulus-related input. The middle ring R2 receives long-range feed-forward input from R1, and the top ring R3 from R2. The couplings from R1 to R2 and R2 to R3 have identical strengths and widths. We deliberately do not introduce any structural feedback coupling to study the capacity of information-theoretical metrics to capture the primitive processing operation of propagating and transferring information through a multi-regional directed hierarchy.

*Reciprocally coupled two-rings network*. We then turn to a second circuit configuration where two regions simultaneously interact via both feed-forward and feed-back connections. Here, the goal is to study the capacity of information-theoretical metrics to track effects of context-dependent top-down modulations (e.g., as in selective attention, Figure 5). In this setup, called 2RC rings (cf. Tables 1 and 2) we reciprocally couple two rings. This closely resembles a model described by Ardid et al. (2007). The bottom ring R1 again constitutes a sensory cortical area, while the top ring R2 represents a prefrontal cortical area. The latter implements working memory, later acting as a source of top-down influences. Parameters of feed-forward and feed-back connections are fine-tuned (see Table 2) to obtain attention-like enhancements of stimulus-response as in Ardid et al. (2007). They are described and commented in detail in the Results section.

### Task simulations

For the one ring configuration of Figure 3 (probing stimulus representation and maintenance), and also for the 3FF rings setup of Figure 4 (probing inter-regional propagation), we generated 1000 trials with different noise realizations per each of four possible orientations *S_pos_* of the stimulus, thus 4000 trials in total, for both the SU and SB working-points. The stimulus injection center positions *S_pos_* are equally spaced along the ring (at angles 0, π/2, π and 3π/2), alternating randomly from trial to trial. Time is measured in arbitrary units *δt* (10 numeric integration steps per *δt*, fourth-order Runge-Kutta integration scheme, augmented with delay). For all analyses, we drop an initial period of 400 *δt* to discard early transients. In each simulated trial, we first record 100 *δt* of baseline dynamics, before injecting a stimulus, which was then maintained for 150 *δt*, i.e. *S*(*t*) = 1 for *t_ON_ =* 100 *δt* < *t < t_OFF_ =* 250 *δt*, and *S*(*t*) = 0 otherwise.

For the analyses shown in Figure 5 (probing working memory and attentional modulation), the task organization is more complex, as there are two stimulus presentations: the first stimulation at position S*_pos_* (“cue” stimulus) started at *t_ON_* = 110 *δt* and stops at *t_OFF_* = 310 *δt*; the second stimulation at positions *S_pos2_* (“match” stimulus) started at *t_ON_*_2_ = 460 *δt* and stops at *t_OFF_*_2_ = 610 *δt.* The cue stimulus position is fixed in all trials at S*_pos_* = π, while *S_pos2_* alternates randomly between 0 and 2π angular positions, in steps of 2π/10. These stimulus combinations were generated for two different conditions, whose rationale is discussed in the Results section. The first condition is called “attention-OFF” (att-OFF), mimicking conditions in empirical experiments in which no attentional modulations are expected (e.g. the “attend OUT” conditions in Treue (2001)). In att-OFF, both the bottom ring R1 and the top ring R2 of the 2RC setup were prepared in the SU state. In this way, bump representations of cue and match stimuli are formed during stimulus presentation, but decay shortly thereafter. The second condition is called “attention-ON” (att-ON), it mimicked experimental conditions in which attentional modulations of response are expected (e.g. the “attend IN” condition of Treue (2001)). The information that attention must be engaged toward the features of the cue stimulus is provided shortly after cue presentation: at time *t_switch_ =* 100 *δt,* the top ring R2 is moved from the SU to a SB regime (parameter modifications are detailed in Table 2). As a result of attention being “switched on”, the top ring R2 is able to maintain a persistent representation of the cue stimulus through the entire delay period between the offset of the cue stimulus at *t_OFF_* and the onset of the match stimulus at *t_ON_*_2_. This maintained working memory representation then interacts nonlinearly with the representation evoked by the match stimulus in R1, producing characteristic gain modulations (see Ardid et al. (2007) and *Results*). To generate Figure 5, we ran 5000 trials for each S*_pos_* and *S_pos2_* combination, in both the attention-ON and the attention-OFF conditions.

### Estimating information-theoretical quantities

In this study we track effects at the algorithmic level of neural function by directly quantifying the way in which simulated circuit dynamic patterns translate into elementary transformations of the information carried by neuronal activity and the presented stimuli. All the information-theoretical metrics we introduce to detect the enactment of different Information Processing Primitives (IPPs) can be seen as elaborations of a few basic quantities (see Cover & Thomas (2006) for a textbook introduction). The amount of information carried on average by observations of a random variable *X* is quantified by Shannon Entropy:

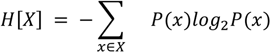

which is a functional of the empirical probabilities *P*(*X*) of observing the different possible values of the variable *X*. The conditional entropy quantifies the amount of information needed to describe the outcome of a random variable *X* given that the value of another random variable *Y* is known, and it is defined as

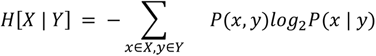

The mutual information (MI) between *X* and *Y* quantifies the statistical dependence between the two variables and it is defined as the difference between marginal and conditional entropies

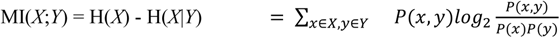

It describes the fraction of information, which is shared, i.e., redundantly encoded by both *X* and *Y*. The Conditional Mutual Information between *X* and *Y* conditioned on a third variable *Z* is defined as:

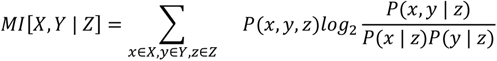

providing the average amount of information carried by *X* (for entropy) or redundantly carried by *X* and *Y* (for mutual information) which is not already carried by Z.

A crucial step to evaluate any information theoretical quantity is the proper estimation of the empirical probability distributions of one or more observables jointly. Here, we used the “plug-in” or “direct” estimators, highly biased for the small amounts of data typically available in neurophysiological experiments, but converging to stable values for large datasets, in which probabilities of different events are estimated as binned frequencies of observation over the collected dataset. We estimate histograms of firing rate variables using 24 equally spaced bins (qualitatively analogous results are obtained using 18 and 32 bins).

For some analyses, when specified in the Results, entropy estimates are computed using a semi-parametric binning-free Gaussian-Copula approach (Ince et al., 2017). In brief, the Gaussian-Copula Mutual Information (GCMI) approach exploits the fact that MI is invariant under monotonic transformations of the marginals. This result can be exploited to render the joint distribution of the variables Gaussian by means of local transformations on the marginals, using the so-called Gaussian copula. GCMI therefore requires transforming the X and Y variables so that the marginal distributions are a standard normal. This copula-normalization involves calculating the inverse standard normal cumulative density function (CDF) value of the empirical CDF value of each sample, separately for each input dimension (i.e., sum-rank computation). Then, entropy values can be estimated using a standard covariance-based formula for Gaussian distributed random variables. We also included a parametric bias-correction for the estimate of the entropy values, which is an analytic correction to compensate for the bias due to the estimation of the covariance matrix from limited data (i.e., limited number of trials). In fact, the limited sampling bias is known to affect the estimation of information theoretical measures (Panzeri and Treves, 1996). The GCMI is a robust rank-based approach allowing to detect any type of relation as long as this relation is monotone. In addition, it can be applied to datasets with a limited number of samples, and with multivariate variables. The GCMI was computed using functions implemented in the Frites Python toolbox (Combrisson et al., 2022). Finally, information-theoretical quantities were normalized by the largest of the entropies of the involved variables (e.g. normalizing MI[*X, Y*] to MI[*X, Y*] / max(H[*X*], H[*Y*])), so that they are bounded in the unit interval and express relative fractions of information rather than absolute amounts.

We now present in detail the specific information-theoretical quantities that we use to define and track Information Processing Primitives (IPPs).

### Tracking Information Processing Primitives (IPPs)

In this study we focused on a simple set of elementary processes of information manipulation stemming from neural activity and use specific information-theoretical functionals in order to detect their emergence and quantify the degree at which different circuit units are engaged into giving rise to them at different times along the simulated tasks. As previously described, the simplest possible operation one can perform with information is to *carry* it. Once a network unit carries some information it can keep carrying it actively for an extended time, i.e. it can *buffer* it; or it can push it to another network unit that did not carry this information before, i.e. it can *transfer* it. The most complicated primitive operation we considered here is *integrating* multiple streams of information, i.e. combining information from multiple sources to reveal the existence of information that was inaccessible to any input source prior to their combination within the integrating node. We now describe in detail these different IPPs and the functionals associated to them.

#### The IPP of carrying information or local encoding of information: Entropy and Mutual Information

The average information carried by the activity of a network unit at a given time is simply given by the functional H[*R*(*t*)], where the variable *R*(*t*) is the across-trials firing rate sampled at time *t*. The probability distribution *P*(*R*(*t*)) is sampled across different trials, all time aligned to the time of stimulus onset (cue stimulus for two ring simulations). One can also evaluate the fraction of the carried information relative to the presented stimulus, either by considering the presence or absence of a stimulus, MI[*R*(*t*), *S*(*t*)], where *S*(*t*) is the spatially inhomogeneous time-course of stimulus presentation, or from the actual feature carried by the stimulus, MI[*R*(*t*), *S_pos_*], where *S_pos_* is the orientation angle of the presented stimulus. Both mutual information terms are then normalized by H[*R*(*t*)] to be expressed in relative form.

#### The IPP of buffering information: Active Information Storage

A circuit node buffering information is a node that keeps carrying some information it was already present at a previous time. Such a primitive processing operation of buffering can be quantified by *Active Information Storage* (Lizier et al., 2012; Wibral et al., 2014). In its simplest manifestation, it corresponds to the mutual information between the past and present activity, MI[*R*(*t*), *R*(*t-τ*)], where *τ* is an adjustable latency, here set to *τ* = 40 *δt*, unless otherwise specified.

Alternatively, one may evaluate the fraction of information about the orientation of the stimulus presented in a trial, which is being actively buffered by a node. The resulting *stimulus-specific active storage* is given by: MI[*R*(*t*), *R*(*t-τ*)] - MI[*R*(*t*), *R*(*t-τ*) | *S_pos_*], i.e., the totally stored information minus the part of this stored information which does *not* depend on the presented stimulus orientation. Note that this measure corresponds to the so-called Interaction Information between the three variables *R*(*t*), *R*(*t*-*τ*) and *S_pos_* (McGill, 1954). Although interaction information can possibly be negative, we always obtain here positive values, as *R*(*t*) and *R*(*t*-*τ*) are generally not independent, and their degree of interdependency does not increase after conditioning on *S_pos_*.

Finally, in all cases, storage measures can as well be normalized by H[*R*(*t-τ*)] to make them relative.

#### The IPP of transferring information: Transfer Entropy

Information flow between neural populations or brain areas can be estimated from the statistical dependencies between neural signals (Bressler and Seth, 2011; Brovelli et al., 2004; Seth et al., 2015) using model-free methods relying on the Wiener-Granger principle (Granger, 1969; Wiener, 1956). It identifies information flow between time series when future values of a given signal can be predicted from the past values of another signal, above and beyond what can be achieved from its autocorrelation. The most general information theoretic measures based on the Wiener-Granger principle are Transfer Entropy (Schreiber, 2000) and Directed Information (Massey, 1990), because they capture any (linear and nonlinear) time-lagged conditional dependence between neural signals (Besserve et al., 2015; Vicente et al., 2011). Transfer Entropy from *X* to *Y* is defined as the conditional mutual information between the past of the *X* and the present of *Y*, conditioned on the past of *Y*

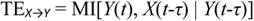

Note that Transfer Entropy is asymmetric, i.e. TE*_X→Y_*≠ TE*_Y→X_,* thus providing a suitable measure for directed functional connectivity (Battaglia et al., 2012; Palmigiano et al., 2017). TE is an information-theoretical generalization of linear Granger Causality (Barnett et al., 2009). We computed two types of transfer entropy. First, active transfer from the stimulus time-course to response rate TE*_S→R_*(*t*) = MI[*R*(*t*), *S*(*t-τ*) | *R*(*t-τ*)] in the one ring configuration of Figure 3. Second, active transfer between the activities *R1_k_* of a unit k in ring *1* and *R2_k_* of another homologous unit (with identical angular coordinate) located in a second ring TE*_R1,k → R2,k_*(*t*) = MI[*R2_k_*(*t*), *R1_k_*(*t-τ*) | *R2_k_*(*t-τ*)], where the first ring is at a lower hierarchical order in the three feed-forward coupled rings configurations of Figure 4. For comparison (and assessment of numerical estimation artifacts), we also compute the backward terms TE*_R→S_*(*t*) and TE*_R2→R1,k_*(*t*) that should be zero by construction since there are no feedback couplings. Again, TE may be normalized by the entropy of the source variable.

#### The IPP of integrating information: Synergistic Modification

A third type of primitive processing operation can arise when two input sources *X*_1_ and *X*_2_ interact and communicate with a common target *Y*. Synergy may then emerge, where extra information is conveyed by the interaction between the sources. This implies that the combined inputs *X*_1_ and *X*_2_ as a whole convey surplus information with respect to the inputs considered separately (Brenner et al., 2000; Latham & Nirenberg, 2005) and the active process of extracting this surplus information –an active process performed by the output node Y– has been called *synergistic modification* (Lizier et al., 2013; 2018).

In the current study, we consider the case where the two source variables are the firing rate of a node in one ring and a stimulus position (*X_1_* and *X_2_*), while the target variable is the firing rate of a node in a second ring (*Y*). We suggest that a key primitive processing operation is the extraction of synergistic information by a target node *Y* from the joint processing of multiple inputs. To do so, we exploit a recent formalism that allows the decomposition of multivariate mutual information between a system of predictors and a target variable, and to quantify the information that several sources (or predictors) variables provide uniquely, redundantly or synergistically about a target variable, the so-called Partial Information Decomposition (PID) framework (Williams and Beer, 2010). The PID formalism can be briefly outlined using the well-known information Venn diagram (see Fig. 5). The total information that the output *Y* carries about the pair of inputs (*X_1_*, *X_2_*) consists of *unique*, *redundant*, and *synergistic* parts. The information *Y* shares with *X_1_*, but not with *X_2_* is commonly denoted as MI(*Y*; *X_1_*\*X_2_*), and, conversely, the other unique information term as MI(*Y*; *X_2_*\*X_1_*). The redundant part is the information provided to *Y* which is shared by both *X_1_* and *X_2_*, denoted MI(*Y*; *X_1_*∩*X_2_*). The remaining part is thus synergy whose amount can be determined by subtracting the non-synergistic fractions of information from the total amount of information MI[*Y*; (*X_1_*, *X_2_*)] that *Y* carries about the pair of inputs. However, these quantities cannot be estimated directly from the data. We chose to operate under the so-called minimum mutual information (MMI) ansatz, which has been shown to provide correct estimations for a broad class of Gaussian systems (Barrett 2015). According to MMI, redundant information can be computed as the minimum of the information provided by each individual source to the target (i.e., one of the two inputs is supposed, as an upper bound estimation, not to carry any unique information)

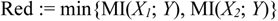

Then, under this ansatz, the synergistic information can be computed by subtracting the mutual information of the single source variables from the total information, adding back the redundancy fraction, otherwise twice subtracted:

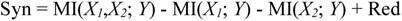

The two equations represent the redundant and synergistic information carried by the co-modulations in firing rates *X_1_* and *X_2_* about the target node *Y*, respectively. It formalizes the process that the output node *Y* may perform in extracting “emergent” information or the above synergistic modification (Lizier et al., 2013). Once again, this metric can be normalized by an entropy term, here the one of the target output *Y*, to evaluate the synergistic fraction of the total output information flow.

## RESULTS

### Two dynamic regimes of response to stimulus

As presented in detail in the *Models and Methods* section, we model a stimulus-selective cortical region as a ring network with a “Mexican-hat” connectivity profile (Figure 2C) (i.e., with more excitatory interactions with units at nearby angular positions along the ring and more inhibitory with nodes farther away). The corresponding phase-diagram is shown in Figure 2D. We measure simulated responses to stimulus presentations (emulating specific tasks) in two different regimes: a *stationary uniform* (SU) and a *stationary bump* (SB) dynamic working point of otherwise similar ring networks (Figure 2D, bottom; see *Models and Methods* for details).

The simplest possible task entails (correctly) responding to the presentation of stimuli with different orientations/angles S_pos_ in different trials. We show simulated recordings of the activity of units in a single receiving region in Figure 3A. The firing rates measured in six exemplar single trials are shown in Figure 3B, three for a region operating in the SU regime (Figure 3B, top) and three for a region operating in the SB regime (Figure 3B, bottom). For each of the trials, we show spatial maps of activity, where the horizontal axis represents time, the vertical axis different units along the ring-network and the firing rate is color coded. The curves below the spatial maps show corresponding firing rate time-series for all units over time. We highlight in color the time-series of units located at specific positions (indicated by matching-color lines on the corresponding spatial maps). As anticipated, we observe a clear distinction between responses in the SU and SB dynamic working points. In the SU regime, a localized stimulus injection generates a bump-like pattern of increased activity emerging around the injection center (red lines and curves) which disappears as soon as the stimulus is switched off. The firing rates of units outside a neighborhood of the injection are either unaffected (blue trace in SU trial #n & trial #j) or, at larger distances from the injection site, may decreases due to the increased lateral inhibition exerted by active units in the bump (blue trace in trial #k). In contrast, in the SB state, bumps of increased activity can develop spontaneously, at random positions along the ring, even without any stimulation, due to the stronger recurrent excitation sufficient to self-sustain reverberation. Upon stimulation, the bump’s position shifts towards the injection position, and remains stable until the end of the stimulus. Finally, after stimulus removal, in the SU state, the stimulus-evoked bumps fade-out while they persist in the SB state.

These two configurations correspond to distinct functional behaviors: stimuli are only *transiently* represented during the SU regime. In contrast, in the SB regime, a sustained activation is observed, supporting the maintenance in *working memory* of the presented stimulus after stimulus removal. The two IPPs concerned with these two simple functional behaviors are associated with “carrying” and “buffering” information, respectively (cf. Figure 1B, top two cartoons), and they can be tracked by different information-theoretical metrics. We focus first on the “carrying” IPP and address the “buffering” IPP in the next section.

### IPP analysis can track the representation of a stimulus

In Figure 3C (top), we show the amount of total information that a ring unit carries as a function of time, averaged over all units and trials, provided by the entropy H[*R*(*t*)] of the activity rates *R*(*t*) as they change as an effect of stimulus presentation. Average entropy of activity is not constant in time and it differs between the SU and SB regimes. In SU, H[*R*(*t*)] is rather low in the absence of stimulus, due to the temporal steadiness and spatial homogeneity of baseline firing rates, fluctuating only due to a weak background noise. Entropy increases, however, during stimulus presentation, as stimulus-evoked bumps emerge and thus produces differences in response rate for different stimulus positions, increasing inter-trial variance. In contrast, in the SB regime, entropy H[*R(t)*] is higher before stimulation, because of the complete stochasticity of the spontaneously emergent bump positions. In our simulated experimental design, stimuli are presented at only four discrete possible positions (a configuration often met also in empirical experiments). Therefore, upon stimulation, the bump positions (and thus the recorded response levels) are quenched to only four pronounced maxima (cf. average response spatial maps in Figure S1A), resulting in a strong entropy reduction with respect to baseline (during which the observed activities vary across trials in a more graded manner). Entropy rises again after stimulus removal, as firing rates are reduced and noisy fluctuations more evident. For both the SU and SB regimes, we furthermore observe tiny peaks and kinks of the Entropy time-courses in correspondence with stimulus onset and offset at times *t_ON_* and *t_OFF_*. These variations can be explained by the non-instantaneous response of the system to instantaneous input variations, causing fast transient dynamics (similar to impulse response) to occur shortly after stimulus onset and offset (so that inter-trial variance is temporarily increased during these transients).

This simple analysis of entropy time-courses is highly specific to the task implemented and reveals no eminent results. Yet, it illustrates how strongly the values of information-theoretical quantities depend on aspects of the neural recordings, such as signal-to-noise ratio and actual task design, which have little to do with algorithmic-level operations. Since entropy is an upper-bound to other metrics (e.g., Mutual Information), absolute entropy variations could result in absolute Mutual Information variations, which do not indicate changes of the way in which information is encoded, but simply reflect changes of the available informational bandwidth. Thus, when attempting to track the manifestations of primitive processing operations, we chose to focus on relative metrics (i.e., normalized by entropy) to evaluate the fractions of the totally available information that are involved in specific processing operations, without having to account for the additional complexity of total entropy fluctuations unrelated to the processing probed.

We then show the relative amount of information that a unit’s activity carries about the stimulus, disentangling two of its aspects: the actual time-course *S(t)* of stimulus presentation, i.e. its presence/absence at specific times and the orientation *S_pos_* of the stimulus, which is a trial-specific property (changed every trial, see *Models and Methods*). This fraction of information is captured by the entropy-normalized mutual information between firing rate *R*(*t*) and time-course of stimulus presentation *S*(*t*) (Figure 3C, middle panel); or stimulus orientation label *S_pos_* (Figure 3C, bottom panel; see Figures S1B-C for spatially-resolved maps of these information theory quantities). Before stimulus presentation, as expected, no information about the stimulus can be extracted from the neural activity, since the entropy at baseline is noise entropy, unrelated to the stimulus. In contrast, during stimulus presentation, activity is modulated in dependence of stimulus position, notably depending on the distance from stimulus angle of the recorded unit.

Starting with information about the stimulus-related inputs time-course *S(t)*, it becomes positive during stimulus presentation, in both the SU and SB regimes, saturating to a higher value in the SU regime (Figure 3C, middle), as the evoked bump activity configuration differs strongly from the homogeneous baseline. During stimulus presentation, some nodes develop exceptionally low or high firing rates which are impossible in spontaneous conditions, thus, signatures of stimulus presence. On the contrary, no *S(t)*-related information exists before or after stimulus presentation. Specifically, information vanishes even after stimulus presentation, and not only before, because, in both cases, spontaneous activity levels in absence of stimulus are equivalent to levels of activity that nodes far away from the site of stimulation injection could have produced even during stimulus presentation (so that it is indistinguishable from spontaneous activity).

Similarly, mutual information with stimulus position *Spos* (Figure 3C, bottom), is absent before stimulus presentations and saturates to a plateau shortly after stimulus onset. It is higher for the SB than the SU regime, as the SB regime provides a larger dynamic range of responses and sharpened bumps. Note that encoding of stimulus is slightly delayed in the SB regime, since the re-arrangement of the self-organized bump positions at baseline is not as fast as the sudden, stimulus-evoked bump injection of the SU regime. In the SU regime, all stimulus-related information vanishes shortly after stimulus offset. In the SB regime, information about stimulus position remains present after stimulus removal, as the stimulus-evoked bump self-sustains itself in its position on the ring. It may slowly drift away under the influence of background noise over timescales longer than the observation window considered here.

Figures S1B-C show that spatial maps of Mutual Information not only depend on time, but also on spatial location. Stripes are clearly visible in these “Infogram surfaces”, because our design comprises only a discrete number of possible stimulus orientations (cf. the stripes in the average spatial map in Fig. S1A). Encoding of stimulus features is in general stronger at locations corresponding to centers of evoked bumps as these locations have the larger dynamic range of variation in our experimental design.

In summary, the normalized Mutual Information of firing rate with stimulus provides an interpretable marker of the IPP of “carrying” stimulus specific information.

### IPP analysis can track the loading and maintenance of a representation in working memory

We are able to detect that the post-stimulus offset activity of units in the SB regime still “carries” stimulus position information. Through which primitive processing operations can this representation be generated and held in working memory? Answering this question requires turning to Information Dynamics metrics, such as Active Information Storage (AIS) (see *Models and Methods*), which quantifies the fraction of the information carried by a node’s activity at a time *t* that was already carried at an earlier time *t – τ*. The process through which this fraction of information is maintained and not discarded corresponds to the IPP of information “buffering” (cf. second cartoon from top in Figure 1B, reproduced in Figure 3D, left).

Figure 3D shows averaged time-traces of Active Information Storage computed with the ring model used in Figures 3A–C. In the SU regime, Active Storage is positive only during stimulus presentation. It reverts to zero after stimulus offset as the stimulus-evoked bumps dissolve back to homogeneously spread baseline activation. In this condition –exactly as in the baseline prior to stimulus onset– all entropy is due to spatially and temporally uncorrelated noise, which is by construction memory-less, thus resulting in null Active Storage.

The situation is different in the SB regime, in which (spontaneous) bump formation is associated with Active Storage and thus positive at baseline and after stimulus offset. However, Active Storage exactly drops at stimulus onset and offset. Indeed, these events induce changes of activity that cannot be predicted based uniquely on prior activity and hence convey information which is not the outcome of “buffering” computations but must come from outside the system. This information injection is instead faithfully captured by another information theoretical metric, Transfer Entropy TE*_S_*_→*R(t)*_ from stimulus to rate (see *Models and Methods*), tracking the complementary IPP of “Transferring”. As shown in Figure 3E, Transfer Entropy peaks match the drops in Active Information Storage visible in the middle panel of Figure 3D. At stimulus onset, network nodes modify their algorithmic role, reducing their implication in the IPP of “buffering” and becoming the recipients of information conveyed by the IPP of “transferring”. Transfer of information from stimulus to activity occurs also at stimulus offset, where a new injection of information indicated by a second peak in Transfer Entropy encodes a release command to either produce bump dispersion (in the SU regime), or a decrease in firing rate together with re-adjustments of the bump shapes (in the SB regime). See also Figure S1D for detailed spatial maps showing the nodes that are most strongly affected by externally injected information at different times.

The information buffered at baseline in the SB regime cannot yet be stimulus specific as the stimulus has not yet been presented. We recall that the positions of spontaneously generated bumps at SB baseline are random. To formalize this intuition, it is possible to quantify the fraction of information about the stimulus, which is stored by network nodes, i.e. the stimulus specific active storage (see *Models and Methods*). The averaged time course of stimulus-specific active storage is shown in the rightmost subpanel of Figure 3D. Its trace correctly captures that the information buffering occurring prior to stimulus presentation is intrinsic, rather than related to a stimulus, while it displays a transient increase after stimulus presentation, as well as during post-stimulus period. Stimulus-specific Active Storage thus provides a valid metric to track the active maintenance of information relative to a presented stimulus.

At the functional level, such stimulus-specific maintenance eventually marks the implementation of working memory. At the algorithmic level, our IPP analyses allow a decomposition of working memory, showing that it arises via the loading of stimulus-specific information –through the IPP of “transferring”– into the activity of the system’s units. These are, by virtue of their collective dynamics, intrinsically devoted to the IPP of “buffering”. This algorithmic decomposition provides not only a narrative of how a system’s dynamics translate into a function, but it also yields a quantitatively precise characterization: suitable information theoretical metrics –Active Storage for “buffering” and Transfer Entropy for “transferring”– provide a precise evaluation of when, where, and how intensively distinct IPPs are performed.

### IPP analysis can track the propagation of representations through a multi-regional hierarchy

We now tackle the algorithmic decomposition of the function of activity propagation: sensory representation propagates through different regions in cortical hierarchy, e.g., from V1 to V2 and above. As detailed in *Models and Methods*, we simulated a feed-forward chain of three ring modules, representing three hierarchically ordered regions (Figure 4A, bottom). The bottom ring R1 represents a sensory cortical area. It receives an input stimulus, which is sent to hierarchically higher cortical areas (R2 and R3). Each unit in the bottom and middle rings (R1 and R2) is coupled to the corresponding unit (and its local neighbors) in the subsequent rings (R2 and R3), respectively.

Figure 4B shows a representative example of single-trial firing rate traces (together with the associated spatial maps of activity) for all three rings, for both the SU (top) and SB regimes (bottom). Red lines and curves indicate units at the position of stimulus injection, blue units far from it, and magenta the initial bump position in SB regime simulations. All panels show the propagation of activity bumps through the hierarchy of rings. Similarly to the case of an isolated ring, bumps are purely stimulus-evoked in the SU regime, while they emerge spontaneously (and are persistent) in the SB regime. The bump maximum amplitude decreases, and its peak latency is delayed when propagating from bottom to top ring. This effect is more pronounced in the SU regime than in the SB regime, where self-amplification via local recurrent excitation acts as a facilitator for propagation. In the SB regime, the effect of forward coupling is already observable without any stimulation: the intrinsic bump positions (magenta lines and curves before stimulus onset) are very similar in the three rings, whereas they would be completely decorrelated if rings were uncoupled.

As for the one ring model shown in Figure 3, we studied whether bump activity performs the basic IPP of “carrying” information about the stimulus. The encoding dynamics revealed by the MI analyses in Figure 4C closely mirror the dynamics of firing rates in Figure 4B. The peak amount of carried information about stimulus position is larger in the bottom ring (black curve) and weaker in the top ring (dotted curve). Furthermore, the rise of encoded information is slower and delayed in rings R2 and R3, particularly in the SB regime where the re-alignment of bump positions is slow and continues in higher order rings even after stimulus offset. Our model thus successfully captures the propagation of sensory representations.

The subsequent question is which IPP is algorithmically mediating this propagation. The obvious and natural candidate to consider is the IPP of “Transferring”, as introduced in the analyses of Figure 3E. In Figure 4D, we show the time series of inter-regional information transfer evaluated via Transfer Entropy. Indeed, we see that Transfer Entropy quickly rises after stimulus presentation, reaching a peak when MI with the stimulus saturates at its maximum plateau value (cf. Figure 4C). Transfer is stronger and faster from R1 to R2 than from R2 to R3, once again in agreement with the description of firing rate dynamics in Figure 4B. After the peak, transfer drops to a plateau level, which slowly decays after stimulus offset. The profile of transfer is more complex for the SB than for the SU regime. Firstly, in the SB state, there is inter-ring transfer of information prior to stimulus onset, since bump positions in R1 and R2 influence the bump positions in R2 and R3, respectively (cf. Figure 4B). Secondly, SB curves are broader and with more marked secondary peaks, associated with the rearrangement of bump positions (and “breathing” of bump widths). The rearrangement of bump positions takes longer than their mere creation. Inspection of the detailed spatial maps of Transfer Entropy shown in Figure S2A shows that substantial transfer occurs even to units far from the stimulus centers as the generation or drift of bumps at locations misaligned with the stimulus must be actively controlled (another inter-regional functional interaction that TE is able to track).

Since the wiring of the considered multi-regional circuit is purely feed-forward, there should not be any significant feed-back information transfer. In the SU regime, we see that backward Transfer Entropy from higher-order toward lower-order rings is close to zero. However, in the SB regime, while feedforward transfer is still larger, allowing to correctly capture the dominant direction of information transfer, a finite, non-vanishing backward transfer is (spuriously) detected. This is due to misestimation of joint probability density given the finite amount of data, as well as to systematic biases of our simple “plug-in” estimators. For instance, we note that the spurious detection of backward transfer is further reduced by using a longer delay in estimating TE (Figure S2B), which allows limiting the impact of fast transients not properly modelled by our quantized estimation of joint activity distributions (see *Methods*). In addition, the use of multivariate delay coordinate embeddings (Takens 1981), as originally prescribed for Transfer Entropy (Schreiber 2000), instead of using just a single delay, suppresses almost completely the inference of spurious backward transfer, as we show here in Figure S2C.

In any case, all estimators, including the simplest ones, were able to correctly detect the existence of dominantly feed-forward information transfer. The “Transferring” IPP is always the main component in the algorithmic decomposition of the propagation of a sensory representation.

### IPP analysis can track the integration of bottom-up and top-down information flows

We move to a last model configuration, specifically designed to reproduce another important cognitive function: selective attention (and the involvement of working memory into its implementation). While attention and working memory are often studied as completely distinct, seminal modeling work by Ardid et al. (2007) has first shown that the attentional effects on sensory responses can be explained as a byproduct of the nonlinear integration of bottom-up inputs from sensory pathways and top-down inputs from a higher-order region. This is in line with earlier hypotheses that working memory could be a fundamental component of mechanisms mediating attentional modulation (Desimone & Duncan, 1995). The latter maintains a working memory copy of a previously presented cueing stimulus with the attended feature value. The attentional effect is depicted as boosting responses to stimuli with attended features and suppressing responses to stimuli with features far from the attended ones (feature-gain-similarity principle, Maunsell & Treue, 2006). We will see that this nonlinear merging of bottom-up and top-down influences can be tracked by the IPP of “integrating”, quantified by synergistic information modification (Figure 5A). In the simple model architecture proposed by Ardid & co-workers (2RC architecture, see *Methods*), two ring networks are reciprocally coupled. The first, lower-order ring R1 represents a generic, selective sensory area tuned in the SU regime, thus generating stimulus-driven bumps. The second, higher-order ring R2, represents the prefrontal cortex, conditionally set to be in SU or SB regime, respectively, depending on attention state “off” or “on”. With att-ON, the second ring becomes thus able to sustain an induced representation of a presented stimulus, even when the stimulus is removed (i.e. it can act as a working memory as in Fig. 3B).

Following Ardid et al. (2007) in the main aspects, we simulate a classic delayed match-to-sample task (Figure 5B). In this virtual task, intended to mimic actual experiments probing the response of cells in MT cortex to drifting random dot patterns, a stimulus is shown within the receptive field of a recorded cell. Cells in MT show a strongly selective response to stimuli drifting in their preferred angular direction (Albright, 1984), resulting typically in bell-shaped tuning curves with a marked unique peak. In our computational model, this selectivity is captured by the heterogeneous responses of units along the lower sensory-area ring, resulting in the response profile given by the black curve in the top panel of Figure 5C. In the virtual task design of Figure 5B, two types of trials exist. A first one, that we call here “att-OFF”, corresponds to a condition in which the standard response of MT cells is measured, in absence of context-specific modulations. Note that, experimentally, this condition is met by having the subject attending actively to features of a second stimulus presented somewhere else. Thus, our “att-OFF” condition matches the “attend OUT” condition of empirical experiments (Martinez-Trujillo & Treue, 2004). The second “att-ON” condition conversely corresponds to the empirical “attend IN” condition where the attentional spotlight is in the receptive field of the recorded cells. The subject is instructed to attend actively to stimuli with the same direction as a first stimulus shown in the cue stage (red stage in Figure 5B). In both conditions, a first cue stimulus is shown and then removed, followed by a delay period of a certain length (light blue stage in Figure 5B) with no stimulus. Then, a second stimulus is presented in the same receptive field, whose direction can be close to or far from the direction of the initially cued stimulus (match stage, orange in Figure 5B). Individual simulated trials for different cue and match stimulus configurations are shown in Figure 5D), in both the attend-ON (Figure 5D, top) and the attend-OFF (Figure 5D, bottom).

When simulating neural responses in the attend-OFF condition (prefrontal area ring R2 tuned in the SU regime), the response profile of the sensory ring to the match stimulus is unchanged with respect to the cue stage. The presentation of match stimuli with different directions simply produces rotated response profiles (black and dashed gray lines in Figure 5C). Inspecting the responses in individual trials, we see that all match stage activity bumps in the sensory ring look similar, and that the activity in the prefrontal ring has a smaller rate and a worse signal-to-noise ratio compared to the second ring (Figure 5E, bottom).

In contrast, in the attend-ON condition the response profile at match stimulus is differently modulated depending on the relative difference of orientation between cue and match stimuli. When the match stimulus has the same orientation as the previously shown cue stimulus (whose copy has been held through the delay period by the second ring, switched to SB), then the response of the direction selective units in the first ring is boosted, while the response of units selective to stimuli far from the attended one is reduced. This can be seen in single trial responses (Figure 5D, top), where the match bump can have darker or lighter hues of gray depending on stimulus configuration. It is even clearer in the red activation profile in Figure 5C, deviating from the black profile for cue and attend-OFF match conditions. Analogous modulations of the response profile, of varying intensities in different locations arise when presenting match stimuli at different directions (orange curves in Figure 5C). The net amount of (simulated) attention-induced modulation can be quantified by computing the percent difference ratio between the response profiles to a stimulus in attend-OFF and ON conditions, which is shown in the bottom panel of Figure 5C. In our virtual task, the positive modulation can be as large as +15% for responses to match stimuli with attended direction and down to between −10% and −40% for match stimuli with unattended direction. Stronger negative modulations occur for stimuli which are ~45° displaced from the attended direction, i.e. roughly corresponding to the lateral width of the attend-OFF tuning curves. We are not going to comment on the model here, as already extensively analyzed in Ardid et al. (2007; 2010), rather, we are going to study the algorithmic effects of its nonlinear dynamics.

We focus specifically on the IPP of “integrating”, occurring here at match stage when the sensory response (*R_1_* in Figure 5D) is the byproduct of nonlinearly merged bottom-up stimulus-related input (S*_pos2_*) and top-down attention-related inputs (*R_2_*). As previously mentioned, the information theoretical quantity we propose to use to track this IPP is synergistic information modification (Lizier et al., 2013; 2014). As graphically depicted in the information Venn diagrams of Figure 5E, the two bottom up S*_pos2_* and top-down *R_2_* inputs carry together (when considered jointly) a certain amount of information MI_tot_ = MI[(*R_2_*, S*_pos2_*); *R_1_*] about what is going to be the output sensory response *R_1_*. A fraction of this total information about the output response is contributed only by one of the two considered inputs. At the exact moment of match stimulus onset, only the bottom-up input S*_pos2_* can convey information about the direction of the newly shown stimulus, while only the top-down input *R_2_*can carry information about the direction of the previously presented cue stimulus. Both these sources of information contribute to determine the final output response and comprise thus two unique information contributions conveyed exclusively by each of the two inputs (unique information fractions, green and blue areas of Figure 5E). In general, some additional information may be shared between the two inputs –including noise entropy not necessarily linked to the task-relevant stimulus– as captured by the redundant information fraction (cyan intersection in Figure 5E). Yet, the sum of unique and redundant contributions could be smaller than the total information MI_tot_. Indeed, some information necessary to determine the response could be conveyed by the two inputs in combination, but by neither of them in isolation. This surplus contribution – “more than the sum of the parts” (Anderson, 1972) – is the synergistic fraction of total information (white area in Figure 5E) and its extraction by the output nodes is termed the synergistic modification operation. We estimate these unique, redundant and synergistic contributions, quantifying specifically when and where along the virtual task of Figure 5B, the activity of the interacting rings implements the emergent IPP of “integrating”.

This extraction of the synergistic surplus of information is tracked and quantified by the information modification surfaces shown in Figure 5F, for both attend-ON (Figure 5F, left) and OFF (Figure 5F, right) conditions (cf. also Figure S3 for more details on the individual terms contributing to its computation). In addition, Figure 5G shows the profile of a section of the modification surface at the beginning of the match stage (averaging range delimited by a dotted black rectangle in Figure 5F). As visible in Figures 5F and 5G, sensory ring units almost do not perform information modification in the attend-OFF condition but do it at specific task-related locations and times in the attend-ON condition.

The most prominent involvement in information modification occurs in the match stage, particularly at the immediate onset of the match stimulus within the dotted black rectangle. This is precisely the stage in which attentional modulations of stimulus response occur. As visible when comparing the profiles of attentional modulation in Figure 5C (bottom) and of information modification in Figure 5G, the participation of a node in information modification is the stronger the more marked the attentional modulations of its activity are. Specifically, information modification is enhanced at the bump flanks, where the most attentional depression is observed. Modification at the bump center position is weaker, because of the weaker overall attentional modulation index and also because the stronger net drive at the bump center helps the recurrent excitatory connectivity within the sensory ring itself to sustain the boosted activity (i.e., the boosting in R1, once “ignited” by an initial amplification trigger signal from R2, can be maintained, in part, locally, and becomes thus less dependent on input integration).

Information modification can be seen to occur even in other parts of the attend-ON activity surface, during different virtual task stages. Modification stripes preceding the match-stage ones can be seen during the delay stage, although with a much weaker intensity. They are related to the fact that some small-intensity activity is observed in the sensory ring during the delay stage, because of some top-down transfer from the working-memory bump in the prefrontal ring (cf. Figure 5F). The nonlinear interactions between the working memory bump and its “sensory shadow” effectively reduce the variability of the sensory ring activity during the delay stage of the attend-ON versus the attend-OFF condition, which is another type of nonlinear phenomenon beyond rate modulations that can result in modification (see *Discussion*). Other modification events occur during the cue stage, probably due to the transient reshaping dynamics of activity bumps following the returning of parameters in the prefrontal ring from SU to SB regime values.

In the attend-OFF state, information modification is much weaker and possibly estimated to small positive values, rather than null, because of numeric estimation artifacts. As detailed by Figure S3, the surfaces shown in Figure 5C are the sum of several other surfaces corresponding to the different terms in the expression for the synergistic information part (see *Methods*). Numeric errors could thus be more important, as more steps are involved.

In conclusion, the function of selective attention admits an algorithmic decomposition involving the IPP of “Integrating”, unlike the simpler functions described in previous sections, decomposing primarily into “Carrying”, “Buffering” or “Transferring” IPPs.

## DISCUSSION

“Information processing” in cognitive sciences is commonly conceived in terms of box-arrow models linking perception to behavior, through multiple stages, not necessarily with explicit reference to neural mechanisms (Fodor, 1968; Rumelhart & McClelland, 1986), but increasingly so (McClelland & Lambón Ralph, 2013), also due to the rise of neuroimaging (Price, 2018). Hypotheses about processing architectures are validated through experimental tasks designed to disentangle the relative contributions and actual relevance of the different boxes in the above box-arrow model. It is difficult, however, to interpret the results of such experiments without implicit reference to categories and concepts of the specific theory being tested (Cooper, 2007). If the postulated theory of how a function works was very different from the –unknown– neuronal computations implementing this function, then the resulting analyses and interpretations would be inherently biased. There is thus a need for data-driven and agnostic approaches to get direct access to the algorithmic level.

Circumventing this epistemological debate, we propose to refer to elementary operations of processing whose implementation can be detected unambiguously in terms of a set of pragmatic metrics applied to the analysis of time series of neural activity. The price to pay for the rigor in the definition of these elementary operations, allowing their quantitative measurement, is that they must necessarily be abstract and act in plain and identifiable ways on raw information conveyed by neural activity. However, the lack of an exclusive relation with cognitive function is compensated by the fact that varying combinations of them can build-up into a variety of different functions. In particular, the primitive operations we considered here –“carrying”, “buffering”, “transferring” or “integrating”– are so low-level and ineluctable that we can hardly think they are not implemented by neural circuits!

Even if these operations are far from the evident functional relevance of computations, such as working memory maintenance or the generation of top-down modulations of activity, they constitute their necessary low-level ingredients, a sort of “neural assembly language”. The analogy with assembly languages is indeed evident in the operation of a conventional digital computer (Wilkes et al., 1951), whose instructions include only the most basic operations, such as erasing or buffering certain memory contents, pushing others to the right or left in a memory register, etc. Despite their deceptive simplicity, these low-level operations are sufficient to give rise to the variety of software outputs that a digital computer can generate, from the word-processors we used to write this article and to the video games and media players that have distracted us during its preparation.

An advantage of focusing on low-level information processing is that such primitive computations can naturally stem from the collective dynamics of a complex non-linear system. For instance, in cellular automata systems, like the famous “game of Life” (https://conwaylife.com/), dynamical patterns known as “gliders” act as self-organized agents of information transfer and their collisions as events of information modification (Lizier, 2013). The complexity of such toy systems is sufficient to endow them with emergent Turing-universal computation capabilities (Adamatzky, 2002). In our study, the IPP of “transferring” materializes by volleys of propagating activity in the coupled rings in Figure 4–emergent dynamic patterns, like gliders in the game of Life–, and the IPP of “integrating” by top-down and bottom-up input activity volleys colliding within the low-level sensory ring in Figure 5. Differently from abstract toy systems, however, the coupled ring models studied here correspond to actual neural circuits mimicking actual cognitive functions. Therefore, the measured information dynamics proceeding from neural dynamics is shown to directly contribute to the modeled functional computations. We can thus fully bridge the gap between the structural level of how the neural circuit is wired (first Marr’s level) and the functional level of which functional computation the neural circuit is aiming at (third Marr’s level). The missing link (the second, under-considered Marr’s level) is provided by the algorithmic decomposition of the simulated computation. Such decomposition indeed precisely quantifies how (through which primitive operations), when (at which epochs within the task), and where (by which network nodes in the multi-regional circuit) information is processed.

Our proposal to seek for primitive processing operations underlying more complex cognitive computations is not completely novel. Training in specific tasks has been shown to automatically confer superior performance in different, apparently unrelated tasks (Singley & Anderson, 1989). This finding led to the speculation that different cognitive algorithms may involve shared processing subroutines and that the acquisition of superior efficiency in these lower-level shared processes would explain the transfer of cognitive skills across tasks. Such notion of “primitive elements” of cognitive processing (Taatgen, 2013) is algorithmic in nature since it refers to manipulations of information which are “pre-functional”, i.e. not necessarily with a simply nameable purpose, but participating in the *implementation* of the final circuit function. Analogously, other cognitive theories postulate the existence of intermediate representations (Wickelgren, 1999; Mel & Fiser, 2000) between the encoding of isolated parts of sensory objects (e.g. a contour segment or a letter sign) and the fully-integrated, context-invariant encoding of whole objects (e.g. a shape or a meaningful word). The generation of such intermediate representations could also be reinterpreted as primitive algorithmic steps towards perception and object recognition. Our notion of IPPs, however, lies at an even lower level than these concepts. Therefore, even pre-functional cognitive operations could still be decomposed into IPPs. This yields a hierarchy of possible algorithmic decompositions, the lowest level given by raw processing directly emanating from coordinated neural activity, quantifiable by information-theoretical functionals such as active storage, transfer entropy and synergistic modification. We are similar in spirit here to previous works such as Ince et al. (2005) or Lungarella & Sporns (2006), extending however their analyses beyond transfer to also encompass buffering and integration.

If the processing operations described by IPP analysis are so unavoidable, one may argue whether the capability to track them really can improve our understanding of neural circuit function. A related discussion can be found in Jonas and Kording (2017) who provocatively asked whether a neuroscientist analyzing recordings of electrical activity within a microprocessor could really understand which computations are performed. Their conclusion was that data-driven analyses alone could at best detect that some interesting processing is ongoing, without hope to really infer its purpose. This is not, however, the scenario we are facing in our study, since we have at our disposal a ground-truth model, knowing both the circuit wiring and the emulated cognitive function. It is precisely the knowledge of ground-truth models that allowed us to realize that different functions are associated with alternative cocktails of IPPs. The spatiotemporal patterns of IPP recruitment we measure in model simulations were reasonably compatible with a priori expectations, thus confirming that IPP analysis yields trustworthy results, successfully able to fingerprint alternate high-level computations, despite its simplicity. The successful proofs-of-concept illustrated by Figures 3–5, allow us to be optimistic about the added-value IPP analyses could bring to real applications.

IPPs go indeed beyond conventional functional connectivity, that just detect which units process information together, by additionally revealing the qualitative type of processing being performed. For instance, in the match stage of the simulated experiment of Figure 5, very similar functional connectivity motifs are generated in attend-ON and OFF conditions, as activity bump configurations in the sensory ring are similarly overlapping in space and time. However, in attend-OFF conditions, functional connectivity motifs primarily capture information *transfer*, while substantial information *modification* is also present in the attend ON condition. The latter cannot be captured by functional connectivity analyses alone. Even without knowing the ground-truth –active merging of top-down and bottom-up influences occurs only in attend ON conditions–, a completely agnostic IPP analysis would have still revealed that two *qualitatively* distinct modes of processing exist, despite only mild *quantitative* differences in activity and activity correlations. As such, we believe that the IPP framework can yield stronger constraints on hypotheses about cognitive processing implementations, guaranteeing that they remain compatible with the complex reality of data. The capacity to track simultaneously different types of processing across different locations will facilitate the identification of putative cognitive architectures combining parallel and sequential aspects (Zylberberg et al., 2011). Indeed, recently, information decomposition techniques could be used to separate large-scale functional interactions between brain regions into synergistic and redundant components, revealing distinct information-processing roles in different cognitive domains (Luppi et al., 2022).

The rate models of neural dynamics considered in our study are extremely simplified with respect to actual, biological neural circuits. Our aim was not to focus on the study of the models –they have been already investigated in depth elsewhere (Ardid et al., 2007)– but to get access to time series of simulated activity from generic neural systems mimicking the performance of actual functions, without bothering about excessive realism for a first proof-of-concept. Despite their simplicity, coupled ring models can nevertheless generate a surprising variety of dynamics. We focused here on asynchronous regimes of activity. Enhancing inhibition or varying delays in the interactions would have given rise to alternative regimes characterized by oscillatory activity, including periodic standing and traveling waves or broadband chaotic oscillations (Roxin et al., 2005; 2006; Battaglia & Hansel, 2011). Ring-models of attentional modulations have been constructed even in the oscillatory regime (Ardid et al., 2010), to uncover the interplay between oscillatory coherence and activity modulations. Simulations in oscillatory regimes could be used, in perspective, to quantify whether the presence of oscillations affects the performed primitive computations (e.g. boosting modification and/or transfer relatively to asynchronous regimes). In addition, the information processing effects of cortical traveling waves (Muller et al., 2018; Chemla et al., 2019) could be elucidated (colliding wavefronts as information modification events?), or to benchmark tools for spectrally-resolved information-theoretical analyses, e.g. frequency-band specific transfer entropy (Pinzuti et al., 2020).

As in our previous studies of state-dependent information transfer by coupled oscillating populations (Battaglia et al., 2012; Palmigiano et al., 2017), we capitalize here on the possibility offered by computational models with carefully shaped structure to generate arbitrarily large quantities of data, in perfectly controlled conditions, which allow a rather straightforward estimation of information-theoretical functionals. Even in this case, estimation is error-prone, as revealed by the spurious inference of information transfer from the higher-to the lower-order rings in the feed-forward configuration of Figure 4. Even if the correct qualitative conclusion is achieved despite misestimation –the dominant direction of transfer is always clearly and correctly identified–, these unwanted results give a warning about the importance to use performing estimators, especially when operating on more limited amount of data as in applications to real data. Here we could still afford using for most analyses a simple binning method for the calculation of entropies and other functionals. However, binning strongly depends on the number of samples, is biased and suffers from the curse of dimensionality (Treves and Panzeri, 1995; Panzeri et al., 2007). In the analyses of Figure S2B-C, we have used a promising alternative based on semi-parametric estimation techniques, namely the Gaussian Copula Mutual Information (GCMI) (Ince et al., 2017). The GCMI has several advantages for large neurophysiological data and brain network analysis: notably, the simplicity of the computation, which renders the algorithms applicable to large datasets with hundreds of variables (e.g., brain regions); and the ability to estimate entropies on few data samples, allowing estimation from short time series containing hundreds of time points, e.g., single trials or across trials. These and other advantages makes GCMI an ideal ingredient for practical toolboxes applicable to real neurophysiological data (Combrisson et al., 2022).

We focused here on a triad of IPPs captured by the quantities of active storage, transfer and modification, including a stimulus-specific version of storage (Figure 3D), which can disentangle information processing relevant to a task from endogenous processing associated to intrinsic dynamics. However, this already rich catalog of IPP metrics is far from being exhaustive. Partial Information Decomposition exists also for more than three variables, yielding a combinatorially-growing number of possible processing types as the number of considered variables increases (Williams and Beer 2010). Furthermore, multivariate frameworks to quantify the informational effects of emergent collective behavior from higher-order interactions have also been proposed for arbitrarily large systems (Rosas et al., 2020). Any additionally defined IPP, independent of its complexity, would still capture informational effects of coordinated dynamics, with relatively more redundant or synergistic styles. The name of “dynome” has been proposed for the collection of possible dynamical modes that a neural circuit with a given connectome can support (Kopell et al., 2014). In our algorithmic-level view, every dynamic pattern within the “dynome” would be seen as an operator on the information conveyed by neural activity. In other words, it would map to an element within an “infome”, or repertoire of information processing modes. Possible examples are transiently coherent oscillatory bursts, serving as information routing enablers (Palmigiano et al., 2017) or the recruitment of distinct system substates, in which the same neurons process information differently at different times (Clawson et al., 2019; Pedreschi et al., 2020). The “storage”, “transfer” and “modification” triad considered here (Figure 1) was sufficient to account for primitive computations that can be performed by one (Figures 3 and 4) or two (Figure 5) interacting activity bumps. However, the list and number of considered IPPs should be tailored to match the variety of intrinsic activity patterns that more general neural circuits engender.

Until now, most attempts to identify canonic computations –such as e.g. inhibition-driven rerouting or normalization (Pouille & Scanziani, 2004; Carandini & Heeger, 2011; Hangya et al., 2014; Miller, 2016)– have strongly committed to specific connectivity motifs being responsible for specific types of processing. Such structure-centric views may be limited by the fact that a connectivity motif can behave differently in different contexts (Aertsen et al., 1989; Nadim et al., 2008, Dahmen et al., 2022), especially when embedded in broader circuits (Kirst et al., 2016), so that the same motif could perform multiple computations. Alternatively, it is possible that very different connectivity organizations implement very similar dynamics (Marder & Goaillard, 2006, Yger et al. 2011, Voges et al., 2012), so that they would give rise to equivalent computations. The quantification of IPPs allows to study information processing directly at the algorithmic level, in a way commensurable with, but “disembodied” from the specific circuit mechanisms producing it. This may allow detecting the action of specific cognitive processes (e.g., attentional modulation) through the identification of their informational signatures (e.g., boosted information modification) even when their effects are more general than a simple rescaling of tuning curves (Helmer et al., 2016). Furthermore, IPP analyses may allow detecting disruptions of primitive information processing itself, in absence of apparent “hardware” damage in the underlying circuits, thus providing a fundamental “software” explanation for widespread cognitive impairments in pathologies (Clawson et al., 2021).

## ACKNOWLEDGMENTS

This work, carried out within the “Institut de Convergence ILCB” (ANR-16-CONV-0002), has benefited from support to NV, DB and AB from the French government, managed by the French National Agency for Research (ANR) and the Excellence Initiative of Aix-Marseille University (A*MIDEX). AB was supported by the PRC project “CausaL” (ANR-18-CE28-0016) and received funding from the European Union’s Horizon 2020 Framework Programme for Research and Innovation under the Specific Grant Agreement No. 945539 (Human Brain Project SGA3). We are also grateful for inspiring discussions with Joe Lizier, Michael Wibral and Stefano Panzeri.

## AUTHOR CONTRIBUTIONS

NV performed all analyses and simulations, JH helped with programming, DB and AB conceived the study, all authors interpreted the results and wrote the study.

## SUPPORTING FIGURES

**Figure S1:**
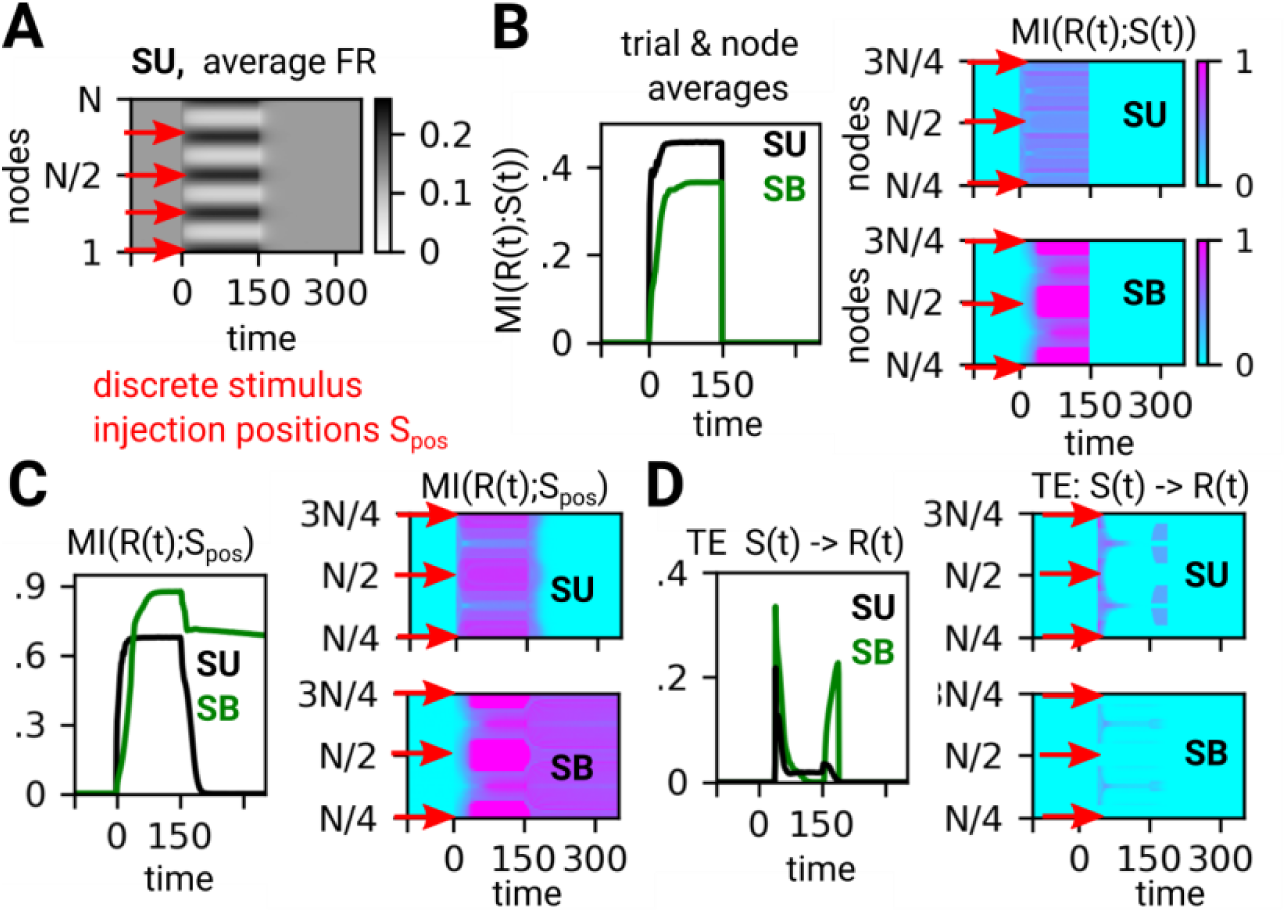
Details about information encoding in the single-region circuit. *(supporting figure to Figure 3)* **(A)** Spatial map (all units vs time) of the trial-averaged firing rates in SU in the single-region model composed of one ring network, red arrows indicate stimulus positions Sp in all trials. This and other spatial maps have a striped appearance due to the use in our numerical experiment setup of a finite, discrete number of possible stimulus orientations, as it is often the case in empirical experiments (stimulus injection positions indicated by red arrows). **(B-D)** Analogous spatial maps for: mutual information between rates and (**B)** stimulus time-course *S(t)* (cf. also Figure 3C middle) or (**C**) stimulus position *S_pos_* (cf. Figure 3C bottom); and (**D**) Transfer Entropy from stimulus time-course to firing rate responses. We note that the stronger transfer from stimulus to rate occurs at the response bump flanks which are the positions experiencing the stronger response modulation (a suppression in this case) as an effect of stimulus presentation.

**Figure S2:**
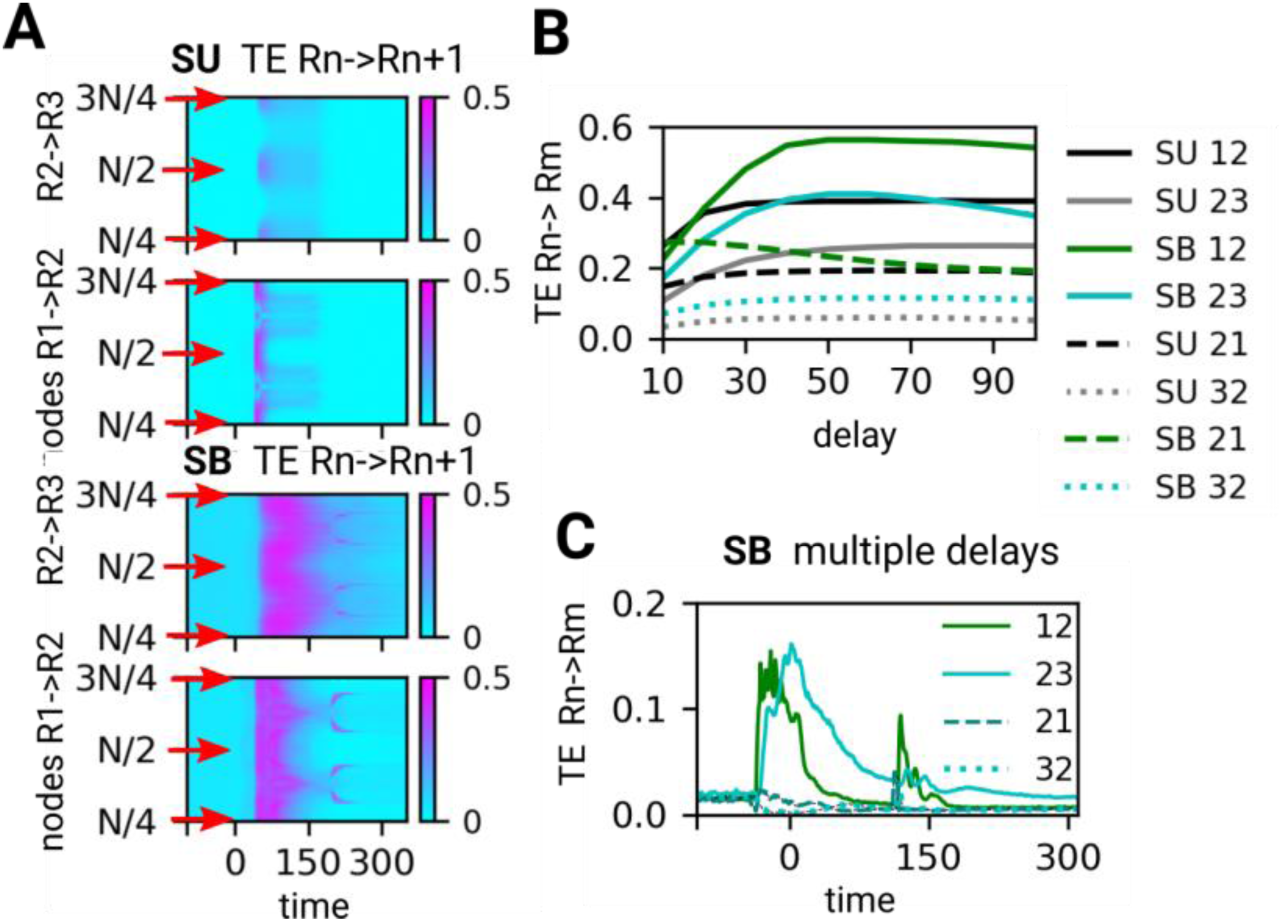
Details on the information transfer in the multi-regional feed-forward circuit. *(supporting figure to Figure 4)* **(A)** Spatial maps (for a subset of units) of the trial-averaged information transfer from ring 1 (R1) to R2 (lower map) and R2 to R3 (upper map). Results from SU state on the top, from SB on the bottom, red arrows indicate stimulus positions S_pos_. **(B)** Dependence of peak transfer entropy TE on the delay used for their calculation (no entropy normalization). Backward transfer is indicated by dotted lines and is reduced for longer delays in the case of R2 to R1 transfer **(C)** Transfer entropy in SB state (trail & node averages) between R1 and R2 and R2 and R3, here calculated through a multivariate scheme using *multiple* delays (dt = 5,10,15,…40 δt) simultaneously (no entropy normalization). See *Methods* for details.

**Figure S3:**
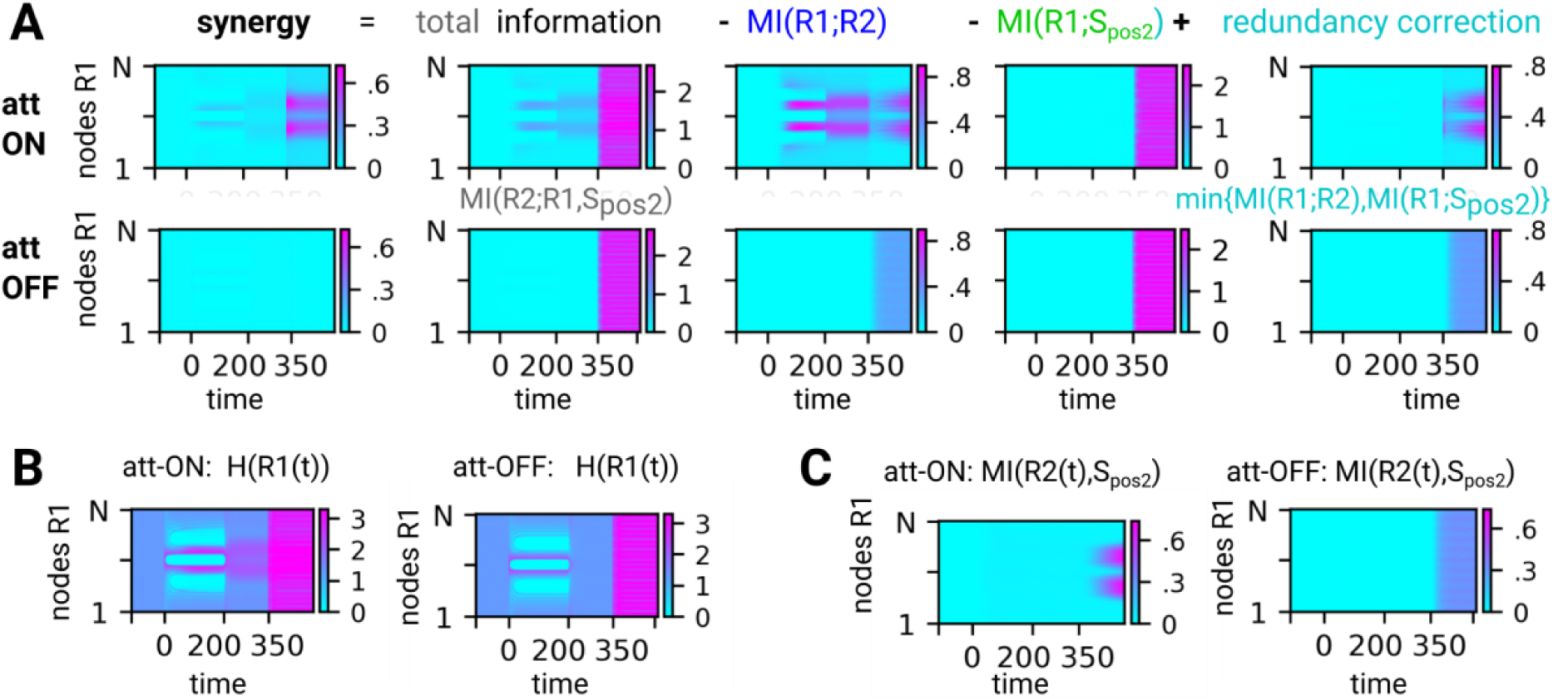
Individual terms of the PID for computing synergistic modification in the bioregional selective attention model. *(supporting figure to Figure 5)* **(A)** Composition of synergy, all the “infograms” for the shown terms are raw and without entropy normalization. Top row, attend ON; bottom row, attend OFF conditions. **(B)** Spatial maps of the entropy of the firing rates in the first sensory ring R1 used for entropy normalizations of the other quantities (left for att-ON, right for att-OFF conditions). **(C)** Spatial maps of the mutual information between the rates in R1 and the stimulus angle shown during match S_pos2_ (left for att ON, right for att OFF). We remark that, to prove that there are no modification effects in the attend OFF condition, the redundancy correction is necessary, as it brings an exact cancelation of twice counted Mutual Information to the stimulus, redundant between the rings.

